# Posterior parietal cortex mediates rarity-induced decision bias and learning under uncertainty

**DOI:** 10.1101/2024.10.25.620383

**Authors:** Weihao Sheng, Xinrui Huang, Yang Xie, Manolis C. Tsakiris, Yang Yang

## Abstract

Optimal decision-making under uncertainty is critical for survival, yet real-world decisions often deviate from optimality. Here, we report a rarity-induced bias in humans and mice, where rare events exert a stronger and more persistent impact on future decisions than common events. Optogenetic manipulations demonstrate two opposing roles of the posterior parietal cortex (PPC): generating rarity-induced bias throughout learning, and driving optimal choices in the expert stage. *In vivo* recordings support both roles: preceding rare rewards enhance the activity of PPC neurons encoding the biased choice while suppressing neurons encoding the opposite choice, causing bias; learning enhances the stimulus-encoding capacity of PPC neurons, promoting optimal choices. A data-driven metacognitive model recapitulated the bias and learning process, and predicted PPC’s causal role in learning, as validated by optogenetics. Our study uncovers an evolutionarily conserved rarity impact in decision-making, and elucidates essential roles of PPC in mediating rarity-induced bias and learning under uncertainty.

## Introduction

In a world that is probabilistic in nature, the ability to learn from uncertain outcomes and make optimal decisions is essential for animals’ survival and well-being^1-3^. Bayesian models provide a normative framework for optimal behaviors in uncertain environments^4^. However, decisions are not always optimal in real life: psychological and economic studies have reported systematic deviation from optimal behavior, in conditions where rare probabilities are involved^5,6^. Specifically, people tend to overweight rare probabilities, acting as if rare events are more likely to occur than they really are^5,6^. This phenomenon of rarity overweighting has significant real-life implications, fundamental to economic behaviors such as gambling and insurance purchasing^7,8^. Psychologists and economists have attributed this behavioral bias to various factors, including the recency, representativeness, and saliency of rare events^5,9,10^, as well as the emotional, psychological, and cognitive states of decision-makers^11-13^.

On top of these factors, a defining feature of any rare event is simply its probabilistic rarity, a purely quantitative property. Surprisingly, this fundamental property inherent in all rare events has been largely overlooked, possibly because rarity is inevitably entangled with other factors in behavioral experiments designed under the *expected utility* or *prospect theory* framework^6,14^. In these paradigms, rarity is often complicated with significant risk, which on its own can influence decisions^12,15,16^, and rare outcomes are typically paired with larger magnitudes of gain or loss^1,3,10,17^, causing rarity to be confounded with saliency. As a result, whether the statistical property of rarity itself, isolated from other factors, has any impact on decision-making remains unknown, and the underlying neural mechanisms have not been explored.

To process the quantitative information of rarity and use it to influence decision-making, the relevant brain regions and neural circuits should be able to: 1) determine event rarity by assessing the probability of recently experienced events from short-term memory; 2) utilize this rarity information to guide future decisions. The posterior parietal cortex (PPC) emerges as a strong candidate for processing the rarity information and implementing it in decision-making: 1) previous work in primates has suggested that PPC plays a critical role in processing numerical information^18^ and probabilities in uncertain contexts^19-21^, and studies in rodents have demonstrated that PPC encodes short-term memories of stimulus, choice, and outcome^22-25^, together supporting PPC’s potential role in assessing probabilistic rarity; 2) PPC is also fundamentally involved in various aspects of decision-making^26-34^, and thus has the potential to influence future decisions.

In this study, we isolated the quantitative property of rarity from other factors to investigate its impact on decision-making and learning, using a probabilistic decision-making task with equal magnitude for rare and common outcomes. We found that rarity has a strong impact on decision-making in both humans and mice, and the rarer the events, the stronger the behavioral impact. Optogenetics show that PPC is required for processing and implementing the rarity impact, supported by *in vivo* physiological recording results that rare rewards bias subsequent choices by bidirectionally modulating the response properties of choice-encoding PPC neurons. In addition, learning dynamically recruits PPC neurons, and monotonically enhances their stimulus-coding capacity at both single-cell and population levels, supporting PPC’s essential role in driving stimulus-based optimal decisions, as also validated by optogenetic experiments. Finally, a data-driven computational model established PPC’s causal role in decision-making and learning under uncertainty.

## Results

### Rare outcomes have a stronger impact on decisions than common outcomes in humans

To isolate rarity from other confounding factors, we designed a probabilistic task with identical rewards for rare (low-probability) and common (high-probability) outcomes, despite the resulting unequal reward expectations (**Figure 1A**). Human subjects were instructed to press the start (S) key to trigger 1 of 4 distinct sounds, and then choose left (L) or right (R) to earn monetary rewards. Two sounds (duck & cat) were associated with 0.8 probability of reward for choosing L (defined as correct choice), and 0.2 probability of reward for choosing R (incorrect choice); reward probabilities were reversed for the other two sounds (canary & frog). Stimuli and common/rare outcomes were pseudo-randomly interleaved (Methods, **Figure S1A**). The outcome was simply shown to the subjects as a money (rewarded, “+”) or no-money (unrewarded, “-”) symbol upon each choice. Most subjects (11/13) eventually reached the expert performance level (defined as 85% correct, **Figure 1B**).

**Figure 1.**
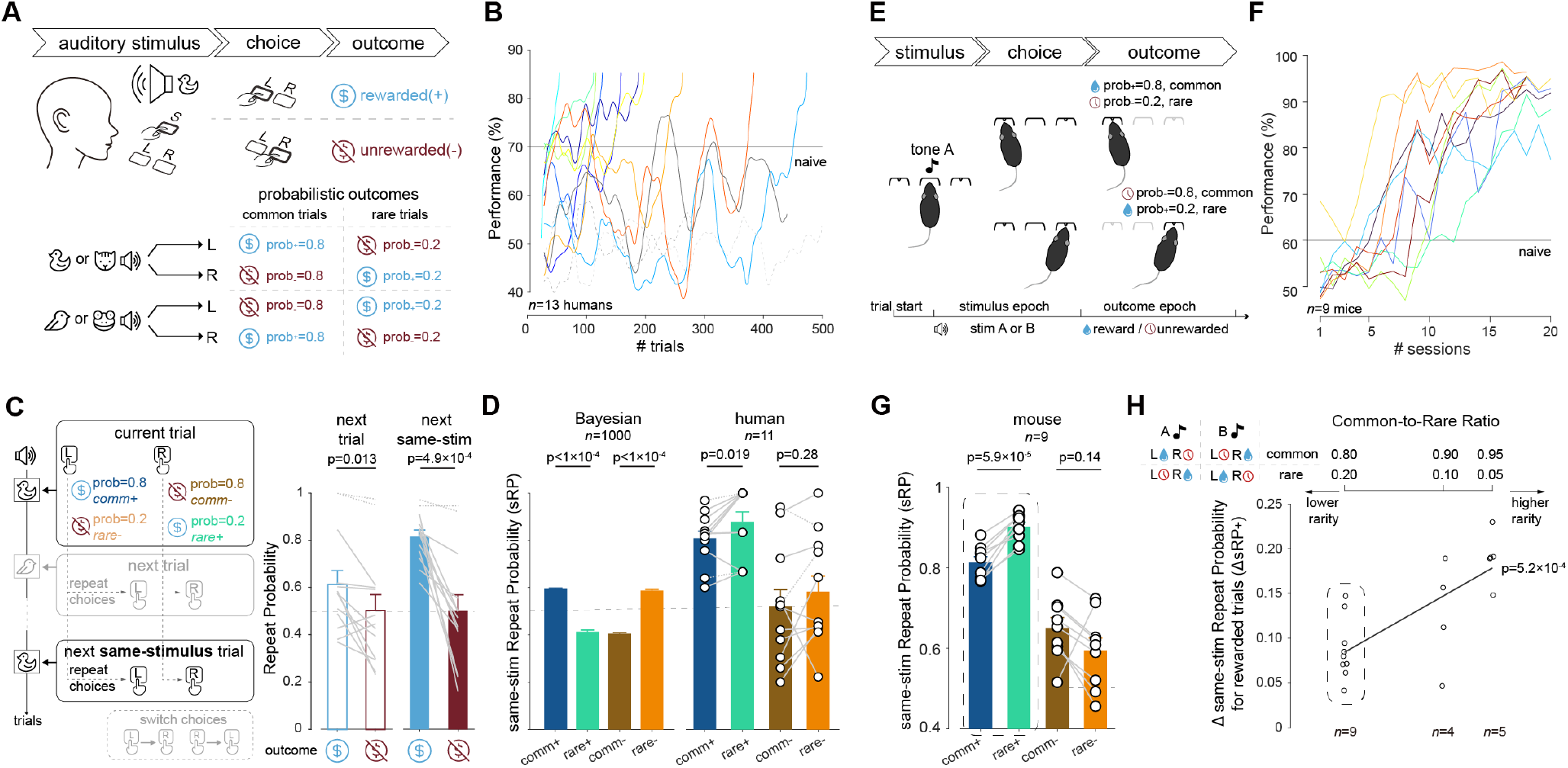
Rare outcomes cause a robust decision bias in humans and mice. **(A)** Human auditory probabilistic decision-making task: for duck/cat sound, L is linked to 0.8 reward probability and R is linked to 0.2 reward probability; probabilities are reversed for canary/frog sound. **(B)** Human subject performance, each line represents one subject. Naïve stage, performance < 70%. **(C)** Left, schematic illustrating choice repeating for next trial and next same-stimulus trial. Right, repeat probability following rewarded & unrewarded trials, for next or next same-stimulus trial, respectively. Wilcoxon signed-rank test. **(D)** Same-stimulus repeat probability (sRP) following different outcomes for Bayesian model and human subjects in the naïve stage. **(E)** Mouse auditory probabilistic 2-alternative-forced choice task. **(F)** Mouse performance across sessions, each line represents one mouse. Naïve stage, performance < 60%. **(G)** sRP following different outcomes in the naïve stage, for mice. **(H)** ΔsRP for rewarded trials (ΔsRP_+_) at 0.8:0.2, 0.9:0.1, and 0.95:0.05 common-to-rare ratios. One-way ANOVA.

As whether an outcome was common or rare was not explicitly shown, subjects need to infer decisions from choices and outcomes of previous trials. Thus, any trial, rare or common, influences subsequent decisions, especially in the naïve stage (defined as performance < 70%, **Figure 1B**), before subjects had learned to make choices based solely on the current stimulus. To evaluate the impact by rare and common trials on subsequent decisions, we computed the choice **R**epeating **P**robability for the **n**ext trial (**nRP**), for both rare and common trials. Given the stimulus-based task design, we also computed the choice **R**epeating **P**robability for the next **s**ame-stimulus trial (**sRP**)^35^ (**Figure 1C**). We observed that sRP, but not nRP, was significantly higher after rewarded trials than after unrewarded trials (**Figure 1C**), indicating a stimulus-based, win-stay lose-switch (WSLS) strategy, which promotes repeating of rewarded choices and switching from unrewarded choices. To simulate the decision-making and learning processes of this task, we built a Bayesian model with equal reward magnitude for rare and common rewards^4^ (Methods, **Figure S1B**). The model predicts significantly higher sRP following rewarded common (*comm*+) choices (*i*.*e*.,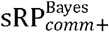) compared to rewarded rare (*rare*+) choices (*i*.*e*., 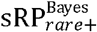) (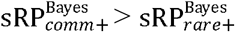, **Figure 1D**), suggesting that subjects will be more likely to repeat choices that have been rewarded more frequently.

However, what we observed in human behavior is just the opposite: subjects are more likely to repeat choices that have been rewarded less frequently (**Figure 1D**). That is, sRP following *rare*+ choices (*i*.*e*.,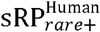) is higher than sRP following *comm*+ choices (*i*.*e*.,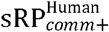) (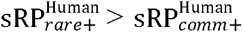, **Figure 1D**). Thus, human behavior apparently violated the Bayesian model in this task. It is worth noting that this is despite the fact that *comm*+ trials occur more often than *rare*+, which should result in enhanced reinforcement of *comm*+ choices, and that repeating a *rare*+ choice leads to a much lower reward expectation than repeating a *comm*+ choice. This surprising result demonstrates a stronger impact on future decisions by *rare*+ than *comm*+, indicating a rarity impact (RI) that strongly biases choices towards *rare*+, causing deviation from Bayesian optimal behavior.

### A mouse model recapitulates the rarity impact

To investigate the neural mechanisms underlying RI observed in humans, we developed a behavioral paradigm in mice, by adding probabilistic outcomes to a classical two-alternative forced choice (2AFC) task^36^ (**Figure 1E**). The mouse pokes into the center port to trigger one of two auditory stimuli (A: 5k Hz pure tone; B: 13 kHz). Stimulus A is associated with 0.8 probability of water reward for left choices and 0.2 probability of the same amount of water for right choices; contingencies reversed for stimulus B. All mice trained in this task showed a sigmoidal learning curve across sessions (300-600 trials in each session; **Figure 1F, Figure S2A**)^37^.

Similar to humans, mice adopted a stimulus-based WSLS strategy (**Figure S2B**). And just like in humans, *rare*+ also had a stronger impact on decisions than *comm*+ in mice, in the naïve stage (performance < 60%, **Figure 1F**), shown by 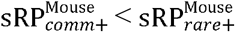 (**Figure 1G**). To investigate whether the strength of RI is associated with the rarity level, we modified the common-to-rare trial ratio to 0.9:0.1 and 0.95:0.05, and trained two separate groups of mice (**Figure 1H, Figure S2A**). We observed larger*Δ*sRP _*+*_ (calculated as: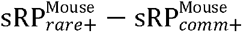) with lower rare ratio (**Figure 1H, Figure S2C, D**), indicating stronger RI for rarer rewards. Therefore, in addition to detecting an evolutionarily conserved rarity-induced bias that leads to suboptimal decisions, we established a behavioral paradigm in mice for studying the neural basis of RI.

### Rarity impact persists across multiple trials

To investigate how the strength of RI changes with time, we took advantage of the fact that sRP_*rare+*_ (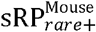 and 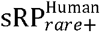) and sRP_*comm+*_ (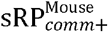 and 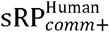) were computed for successive same-stimulus trials separated by different numbers of other-stimulus trials in between. This allowed us to assess how sRP_*rare+*_ and sRP_*comm+*_ evolved as the in-between trial number increased (**Figure 2A**). By computing sRP_*rare+*_ and sRP_*comm+*_ for different in-between trial numbers, we found that as in-between trial number increased, sRP_*comm+*_ decayed quickly while sRP_*rare+*_ remained stable (**Figure 2A, B**), in both humans and mice. These results indicate that the impact of *rare*+ trials on future decisions decays more slowly with time than the impact of *comm*+ trials. For unrewarded trials, the same trend was observed in mice (**Figure 2B**): the impact of *rare*- trials also decays more slowly than that of *comm*- trials. These results demonstrate that rare outcomes have a long-lasting impact on decisions, resilient to distraction and memory decay.

**Figure 2.**
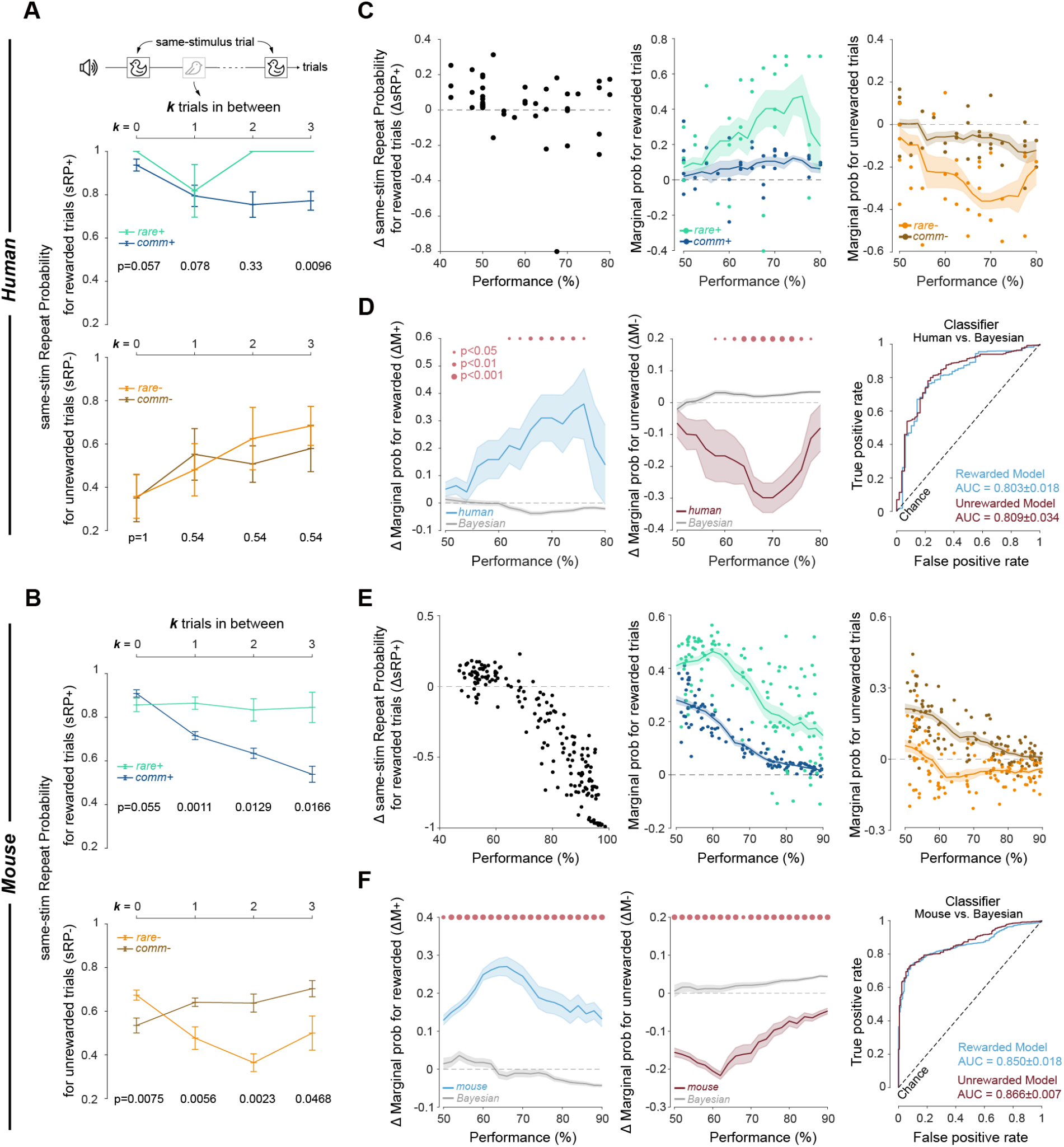
Rarity impact persists across multiple trials and throughout learning. **(A)** sRP_+_ (top panel) and sRP_-_ (bottom panel) for different numbers (***k***) of other-stimulus trials in between two adjacent same-stimulus trials, for humans. Wilcoxon rank sum test. **(B)** Same as (**A**), for mice. **(C)** Left, ΔsRP_+_ across performance for humans, each point representing a 40-trial block. Middle & right, marginal probabilities for *rare+* (M*_rare+_*), *comm+* (M*_comm+_*) & *rare-* (M*_rare-_*), *comm-* (M*_comm-_*) across performance. **(D)** Left & middle, ΔM_+_ (M*_rare+_*-M*_comm+_*) and ΔM_-_ (M*_rare-_*-M*_comm-_*) for humans and Bayesian model. Red dots indicate significant difference between human behavior and Bayesian model, significance level represented by the dot size. Right, receiver operating characteristic (ROC) curves for classifiers predicting human vs. Bayesian model (n=11, 5-fold cross-validation). **(E)** Same as (**C**), for mice. Each point represents one session. **(F)** Same as (**D**), for mice (n=9).

### Rarity impact persists across learning

Another informative trend we observed was that as performance improves, ΔsRP_*+*_ decreases (**Figure 2C, E**, left). While ΔsRP_*+*_ *>* 0 obviously indicates the presence of RI, it is important to note that ΔsRP_*+*_ *<* 0 does not necessarily mean the absence of RI. This is because, in addition to the WSLS strategy through which RI manifests, decisions are also influenced by the current stimulus. As learning progresses and the behavioral performance improves, more choices were made optimally, *i*.*e*., following the stimulus-based optimal strategy. This strategy pushes choices towards *comm*+ (optimal) choices and away from *rare*+ (suboptimal) choices, resulting in increased sRP_*comm*+_ and decreased sRP_*rare*+_.

To assess RI without the confound of the stimulus-based strategy, we define marginal probabilities, M_*rare*+_ and M_*comm*+_, as a measure of the impact of *rare*+ and *comm*+ on subsequent decisions, respectively (Methods). To compute M_*rare*+_, we calculated the difference between the conditional probability of making a *rare*+ choice (*i*.*e*., incorrect choice) following a *rare*+ trial, and the probability of making a *rare*+ choice following any trial:

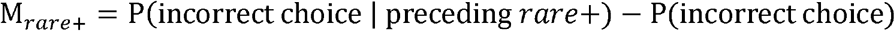

Note that P(incorrect choice | preceding *rare*+) equals sRP_*rare+*_, and P(incorrect choice) equals (1 <Xpx class=“byline”>-performance), performance being the proportion of correct choices in the block (for humans, **Figure 2C**) or the session (for mice, **Figure 2E**). Similarly,

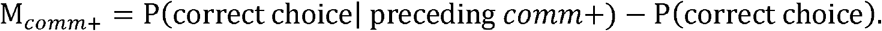

RI can be measured as the additional impact of *rare*+ trials on future decisions, compared to the impact by *comm*+ trials on future decisions 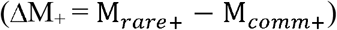.In the Bayesian model, 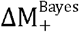 (calculated as 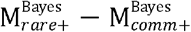) is close to 0 (**Figure 2D, F, Figure S2E**), which is expected because the Bayesian model does not have RI. In humans and mice, however, 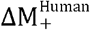 and 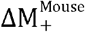 are consistently above 0 and significantly different from 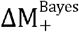,across all performance levels (**Figure 2D, F**)^38^, indicating that RI persists even when most decisions are optimally made. Interestingly, although the additional influence by *rare*- trials on future decisions compared to *comm*- trials was not detectable by computing sRP_*rare*-_ and sRP_*comm*-_, we found 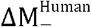 and 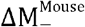 (calculated as 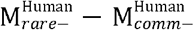,and 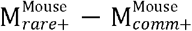,respectively) to be consistently below 0, and significantly deviate from the Bayesian model (**Figure 2D, F, Figure S2E**), demonstrating the presence of RI for unrewarded trials. Thus, rarity exerts a significant impact on decision-making throughout task performance, and such impact prevails regardless of the presence of rewards.

### PPC neural activity is necessary for generating rarity impact

Previous studies have shown that the posterior parietal cortex (PPC) encodes probability information^19-21^ and short-term behavioral history^22-25^, both critical for RI processing. To investigate whether PPC is required for RI, we used optogenetics (eNpHR3.0) to bilaterally inhibit PPC, during either the stimulus epoch (*stim-opto*) or the outcome epoch (*outcome-opto*) of rare trials (**Figure 3A-C, Figure S3A**). A masking yellow light was delivered in the behavioral chamber in all trials to ensure that mice did not use the stimulating light as an additional cue. We found that while RI was unaffected in *outcome-opto* mice, it was significantly reduced in *stim-opto* mice, as shown by significantly lower ΔsRP_+_ and higher ΔsRP_-_ compared to control mice (**Figure 3D, Figure S3B**). Consistent with these results, in *stim-opto* mice, *rare*+ trials no longer had a persistently stronger impact on future decisions than *comm*+ trials (**Figure 3E**). In addition, significant reduction of RI in *stim-opto* mice was observed throughout learning, as measured by ΔsRP and ΔM across performance levels, for both rewarded and unrewarded trials (**Figure 3F-H, Figure S3C**). These results establish that PPC activity is required for generating RI, specifically in the stimulus epoch of rare trials.

**Figure 3.**
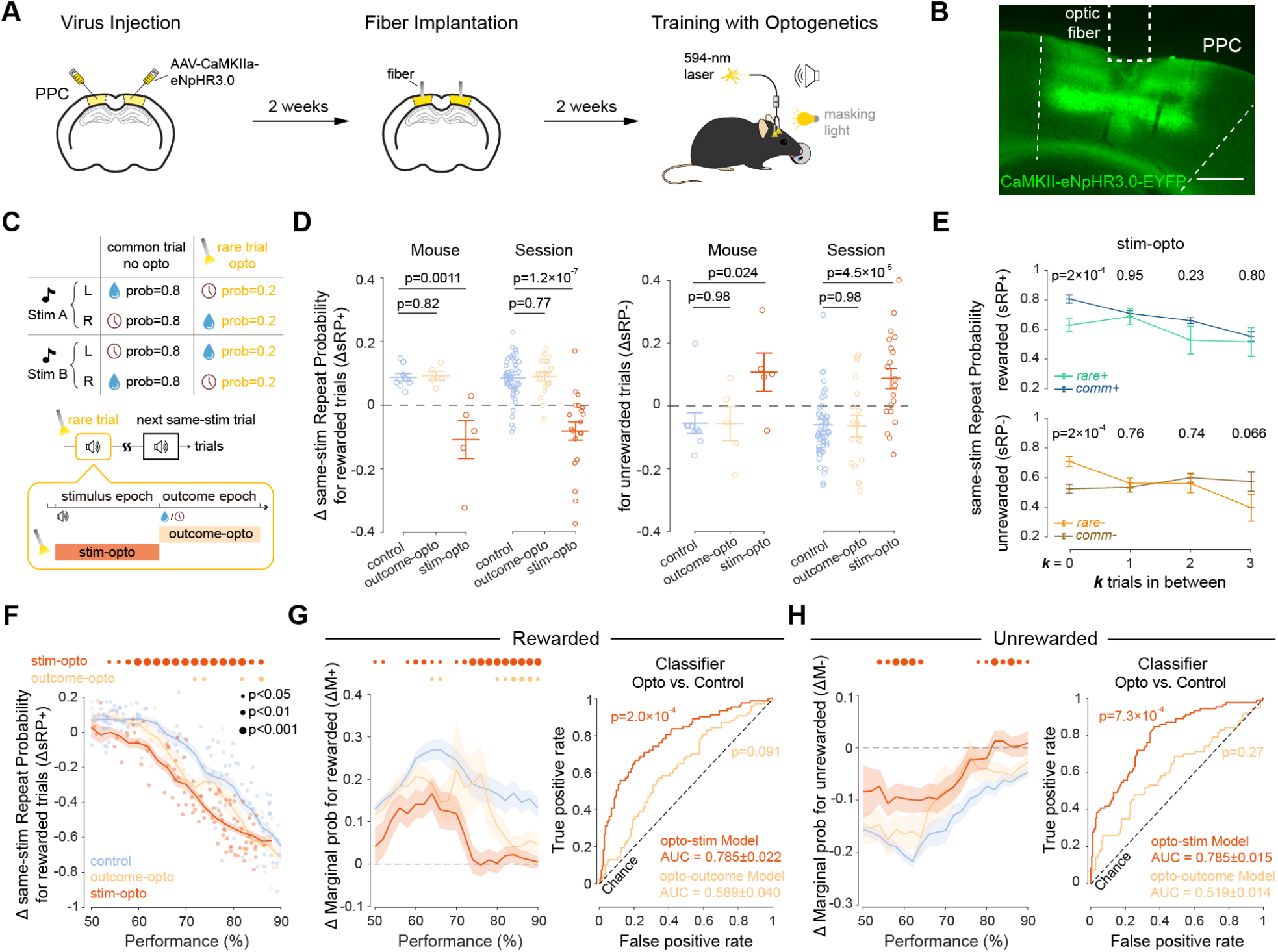
Optogenetic inhibition of PPC reduces the rarity impact. **(A)** Timeline for optogenetics experiments. **(B)** Histology image showing the virus expression and the optogenetic inhibition site. **(C)** Schematic of the optogenetics experiments: yellow light was delivered either during the stimulus epoch (*stim-opto*) or outcome epoch (*outcome-opto*) of rare trials. **(D)** ΔsRP*_+_* (left) and ΔsRP*_-_*(right) for control, *outcome-opto* and *stim-opto* mice. Wilcoxon ranksum test. **(E)** sRP_+_ (top panel) and sRP_-_ (bottom panel) for different numbers (***k***) of other-stimulus trials in between two adjacent same-stimulus trials, for *opto-stim* mice. Wilcoxon signed rank test. **(F)** ΔsRP*_+_* across performance levels for control, *stim-opto*, and *outcome-opto* mice. Wilcoxon ranksum test. Red and orange dots indicate significant difference between optogenetics and control mice, significance level represented by the dot size. Bayesian model. **(G)** Left, ΔM_+_ (for rewarded trials) across performance for control, *stim-opto*, and *outcome-opto* mice. Red and orange dots indicate significant difference between optogenetics and control mice, significance level represented by the dot size. Right, ROC curves for classifiers predicting *stim-opto* vs. control, and *outcome-opto* vs. control for rewarded trials. (**G**) Same as (**F**), for unrewarded trials.

### PPC encodes rarity information

PPC is involved in processing mathematical information, including probability^18-21^. To investigate how PPC neurons encode probabilistic rarity, we made extracellular recordings in mouse PPC during task performance (5 mice, 100 behavioral sessions, **Figure 4A, Figure S4A-F**), and isolated 3135 single units. We explored rarity encoding by examining how rare trials modulate PPC neuronal activities in subsequent trials. To do this, we categorized trials into 4 groups based on the preceding trial outcome (**Figure 4A**, bottom). Focusing first on trials following *rare*+ (**Figure 4A, B**), we used receiver operating characteristic (ROC) analysis to calculate a *rare*+ modulation index (RMI_+_), which measures the difference in a neuron’s firing rate in trials following *rare*+ trials compared to its firing rate in trials following other trials (Methods). We observed that ∼20% of recorded PPC neurons were significantly modulated by *rare*+, with roughly half exhibiting enhanced activity and half suppressed activity (**Figure 4B, C**). Consistent with the persistence of RI in behavior throughout learning, we also observed *rare*+ modulated PPC neurons across all performance levels (**Figure 4D**). Moreover, the strength of RI, as measured by ΔM_+_, was positively correlated with the proportion of PPC neurons that were significantly modulated by *rare*+ (**Figure 4E**). Similarly, we also observed *rare*- modulated PPC neurons across performance levels, and ΔM_-_ was negatively correlated with the proportion of PPC neurons modulated by *rare*- (**Figure S5A-F**). These results indicate that PPC encodes probabilistic rarity through modulation of neuronal activity by preceding rare trials.

**Figure 4.**
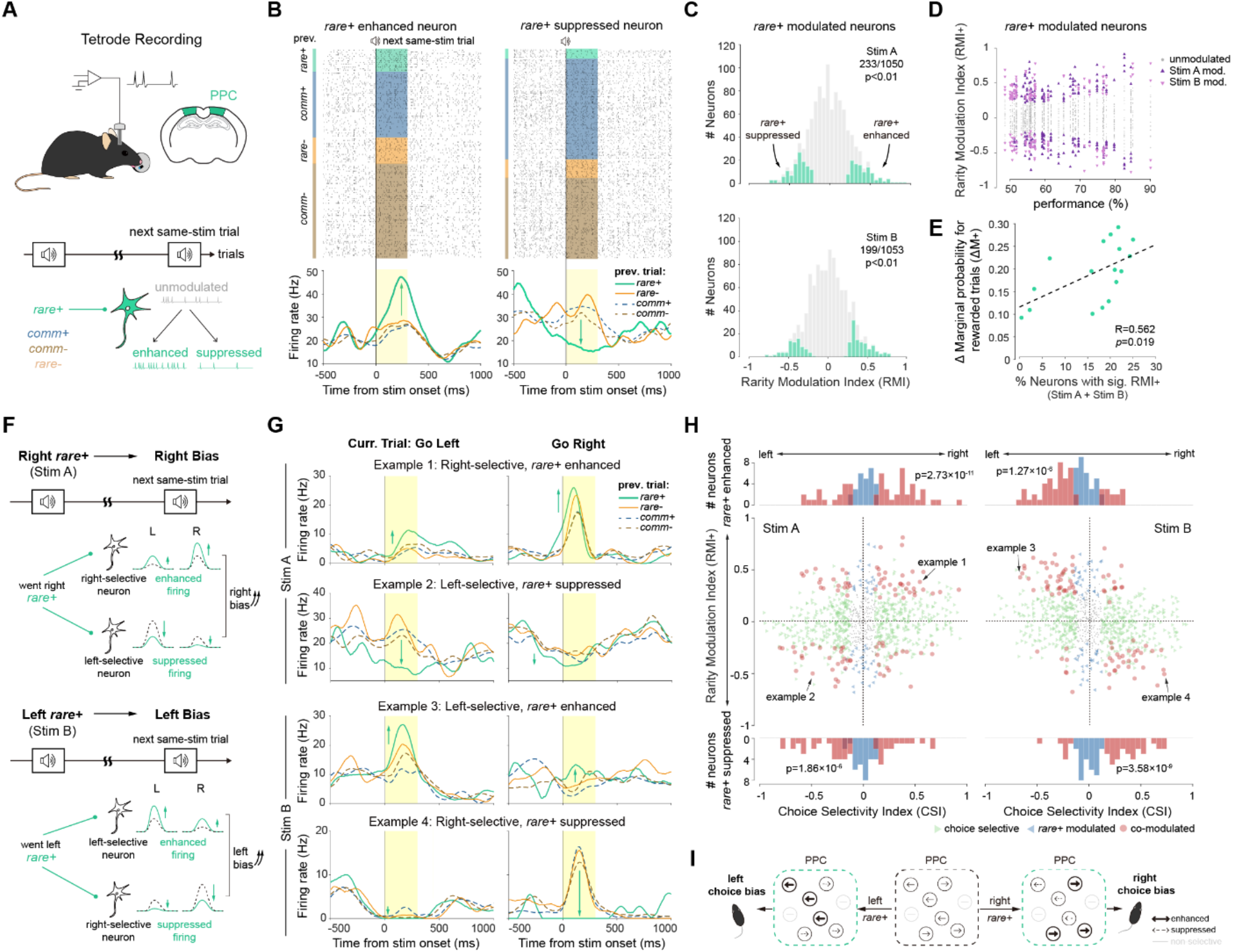
Rare rewards modulate choice-encoding PPC neurons to induce decision bias. **(A)** Schematic showing tetrode recording in PPC, and modulation by preceding *rare*+ trials on neuronal activity. **(B)** Spike raster and peri-stimulus time histogram (PSTH) of 2 example neurons, showing enhanced (left) or suppressed (right) activity following *rare*+ trials. **(C)** Histogram showing *rare*+ modulation index (RMI_+_) of PPC neurons calculated by ROC, for stimulus A (top panel) and stimulus B (bottom panel). Color bars represent significant modulation by *rare*+ (bootstrapping). **(D)** RMI_+_ of individual neurons recorded across performance. Triangles, neurons significantly modulated by stimulus A or stimulus B *rare*+. **(E)** Correlation between the RI strength (ΔM_+_) and the fraction of neurons with significant RMI_+_. **(F)** Top, schematic showing preceding right *rare*+ enhances the activity of a right-preferred PPC neuron, and suppresses a left-preferred PPC neuron. Bottom, schematic showing left *rare*+. **(G)** PSTH of 4 example choice-encoding neurons modulated by *rare*+, in left-going and right-going trials. Example 1, a right-selective neuron enhanced by preceding right *rare*+ in both left- and right-going trials. Example 2-4, other modulation types. **(H)** RMI_+_ plotted against choice selectivity index (CSI) for all neurons. Top panel, stim A; bottom panel, stim B. Each point represents one neuron. Red points, neurons significantly co-modulated by current choice and preceding *rare*+. Histograms show CSI for *rare*+ enhanced (top) and *rare*+ suppressed (bottom) neurons. Kolmogorov-Smirnov test. **(I)** Schematic demonstrating *rare*+ modulation of choice-encoding neurons and choice bias.

### Rarity bidirectionally modulates choice-encoding PPC neurons to influence choices

Previous studies have shown that PPC activity can directly guide behavioral choices^26,28,32-34^. Thus, it is plausible that preceding rare trials modulate choice-encoding PPC neurons to influence choices (**Figure 4F**). To investigate this, we first examined the choice selectivity of PPC neurons during the stimulus epoch, which precedes the animal’s choice behavior (Go left/Go right, **Figure 4G**). Using ROC analysis, we computed a choice selectivity index (CSI) for each neuron. We found that most PPC neurons fired significantly more for either right choices (termed “right-selective neurons”) or left choices (“left-selective neurons”) (**Figure S6A-C**). Increased firing rates in right-selective neurons are associated with increased right choice probability and decreased left choice probability; the opposite is true for left-selective neurons. Therefore, a plausible mechanism for how rarity modulation of PPC induces choice bias, is that preceding rare trials can increase the firing rate of PPC neurons selective for the biased choice, while suppressing neurons selective for the opposite choice (**Figure 4F**).

This is indeed what we found. Recall that RI is evident as preceding right-going *rare*+ trials (*i*.*e*., *rare*+ of stimulus A) led to a right bias by increasing right choice probability; similarly, left-going *rare*+ led to a left bias (**Figure S7A**). Focusing first on right *rare*+, we found that right-selective neurons were enhanced in both left- and right-going trials following right *rare*+ (example 1, **Figure 4G, H**), leading to increased right choice probability (**Figure 4I**), regardless of the choice motion. Conversely, left-selective neurons were suppressed by right *rare*+ (example 2, **Figure 4G, H**), also contributing to the right bias (**Figure 4I**). Likewise, preceding left *rare*+ enhanced the activity of left-selective neurons and suppressed right-selective neurons (examples 3 & 4, **Figure 4G, H**), leading to a left bias (**Figure 4I**). This influence is independent of reward, as combining *rare*+ and *rare*- produces the same effect (**Figure S7B**). These results reveal the neural mechanism that rarity biases subsequent decisions by bidirectionally modulating the activity of choice-encoding PPC neurons.

### Learning under uncertainty enhances stimulus encoding of PPC neurons to drive decisions

Despite the persistence of RI, most human subjects and all mice eventually learned to make stimulus-based decisions. While PPC is known to be involved in stimulus-based decision-making and learning^26,28-34,39,40^, it remains unclear how PPC participates in learning when the outcomes are uncertain. To address this, we used a two-way ANOVA considering stimulus and choice^26,28,32-34^, to identify stimulus-encoding, choice-encoding and dual-encoding PPC neurons across learning. The proportion of choice-encoding neurons remained high throughout task performance, but the proportion of stimulus-encoding neurons changed dynamically, showing a sharp increase around 65% performance level and a gradual decline after 85% performance level (**Figure 5A**). These results suggest that learning dynamically recruits PPC neurons for stimulus encoding, particularly when performance is quickly improving. To further evaluate the strength of choice and stimulus encoding across learning, we computed choice and stimulus discriminability (d’) for each neuron^41^. Choice d’ remained high (**Figure S8A, B)**, while stimulus d’ increased significantly as performance improved (**Figure 5B, Figure S8C**). Using demixed PCA^42^ to assess population encoding of choice and stimulus, we observed extremely low population encoding for the stimulus at performance below 65%, followed by a continuous increase (**Figure 5C**). Thus, stimulus-based learning under uncertainty is accompanied by an increasing stimulus-encoding capacity of PPC, at both single-cell and population levels (**Figure 5D**).

**Figure 5.**
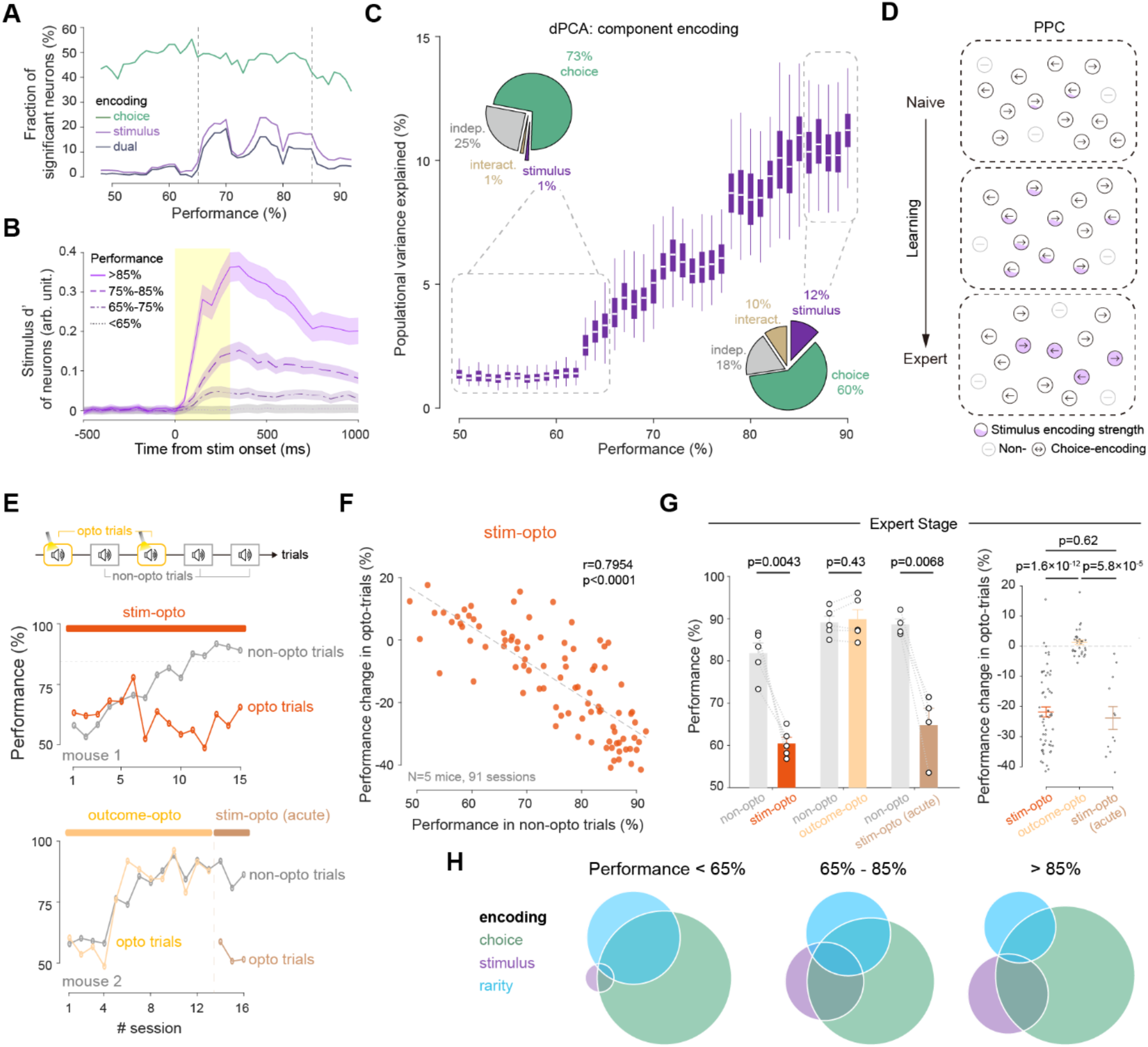
Stimulus-guided learning under uncertainty enhances stimulus encoding of PPC neurons to drive decisions. **(A)** Fraction of significant choice-encoding, stimulus-encoding, and dual-encoding neurons across performance. Two-way ANOVA (p<0.01). **(B)** Average stimulus discriminability (d’) for stimulus-encoding neurons across performance ranges. **(C)** Population variance explained by stimulus, calculated using demixed PCA with 200 neurons (bootstrapped 100 times) recorded in sessions within the performance range of [-5%, +5%]. Boxes show 25^th^ and 75^th^ percentiles. Insets, population variance calculated using dPCA with neurons recorded at <65% or >85% performance. **(D)** Schematic showing strength of stimulus encoding and fraction of neurons in PPC throughout learning. **(E)** Performance of an example *stim-opto* and *outcome-opto* mouse in *opto* and *non-opto* trials across sessions. The *outcome-opto* mouse was later switched to *stim-opto(acute)* after reaching 85% performance. **(F)** Difference in performance between *stim-opto* and *non-opto* trials throughout learning in each session. **(G)** Performance in *non-opto* and *opto* trials at expert stage for each mouse (left, Wilcoxon signed-rank test) or each session (right, Wilcoxon ranksum test). **(H)** Overlap of choice-encoding, stimulus-encoding, and *rare*+ modulated neurons at <65% (neuron number: 528, 24, 255, respectively), 65∼85% (318, 121, 141), and >85% (15, 5, 4) performance levels. The neuron number for >85% performance is very small because for a neuron to be included in the analysis, the behavioral session in which it was recorded had to have more than 3 *rare*+ trials. But at expert stage mice rarely make *rare*+ (suboptimal) choices.

To examine whether the stimulus-encoding capacity of PPC neurons is causally linked to decision-making, we assessed mouse performance while bilaterally inhibiting PPC in a subset of trials (**Figure 5E**). Consistent with the increasing stimulus-encoding capacity during learning, the performance impairment in PPC-inhibited trials increased as learning progresses (**Figure 5E, F**), becoming highly significant at expert performance levels (**Figure 5G**). To ensure that mice were not using the stimulating light as an additional cue, we switched the inhibition window to

the stimulus epoch after a group of *outcome-opto* mice reached expert levels (*stim-opto (acute)*, **Figure 5E**). We observed an immediate performance impairment after the inhibition window switch (**Figure 5E, G**). These findings demonstrate that as learning progresses, the strength of stimulus encoding by PPC increases, and such stimulus encoding contributes to stimulus-based decision-making. Notably, the stimulus-encoding and rarity-modulated PPC neurons are partially overlapping (**Figure 5H**), suggesting that the two opposing roles of PPC in behavior involve overlapping neuronal populations.

### A metacognitive model reveals PPC’s causal impact on learning under uncertainty

We have demonstrated PPC’s two opposing functions in decision-making: mediating a rarity-induced bias, and driving stimulus-based decisions. These two functions can influence learning in opposite directions: the former impedes learning, while the latter promotes learning. To gain insights into the learning process, we aimed to simulate our data with learning models. We tested 3 established computational models: a Bayesian model^4^, a WSLS model^43^, and a reinforcement learning (RL) model^44^. Although we used a wide range of parameters, none of the models recapitulated the rarity-induced decision bias (**Figure S9A-F**), thus unable to accurately capture the learning behavior in this task.

To address this question, we developed a data-driven dual-agent model, with each agent representing a behavioral impact encoded by PPC (Methods, **Figure 6A, Figure S10A**). The first agent implements rarity-based WSLS (r-WSLS, **Figure S10B, C**)^43^, in which WSLS probabilities are updated trial-by-trial based on an estimate of the rarity state^2,45,46^. The second agent implements RL^44^, using a multiplicative update rule^37^ and different learning rates for rewarded and unrewarded trials^47^. To integrate the two agents, we implemented a metacognitive framework that combines them using normalized decision weights derived from entropy-based metrics in the RL agent^48^, which is the primary driver of learning (Methods, **Figure 6A**).

**Figure 6.**
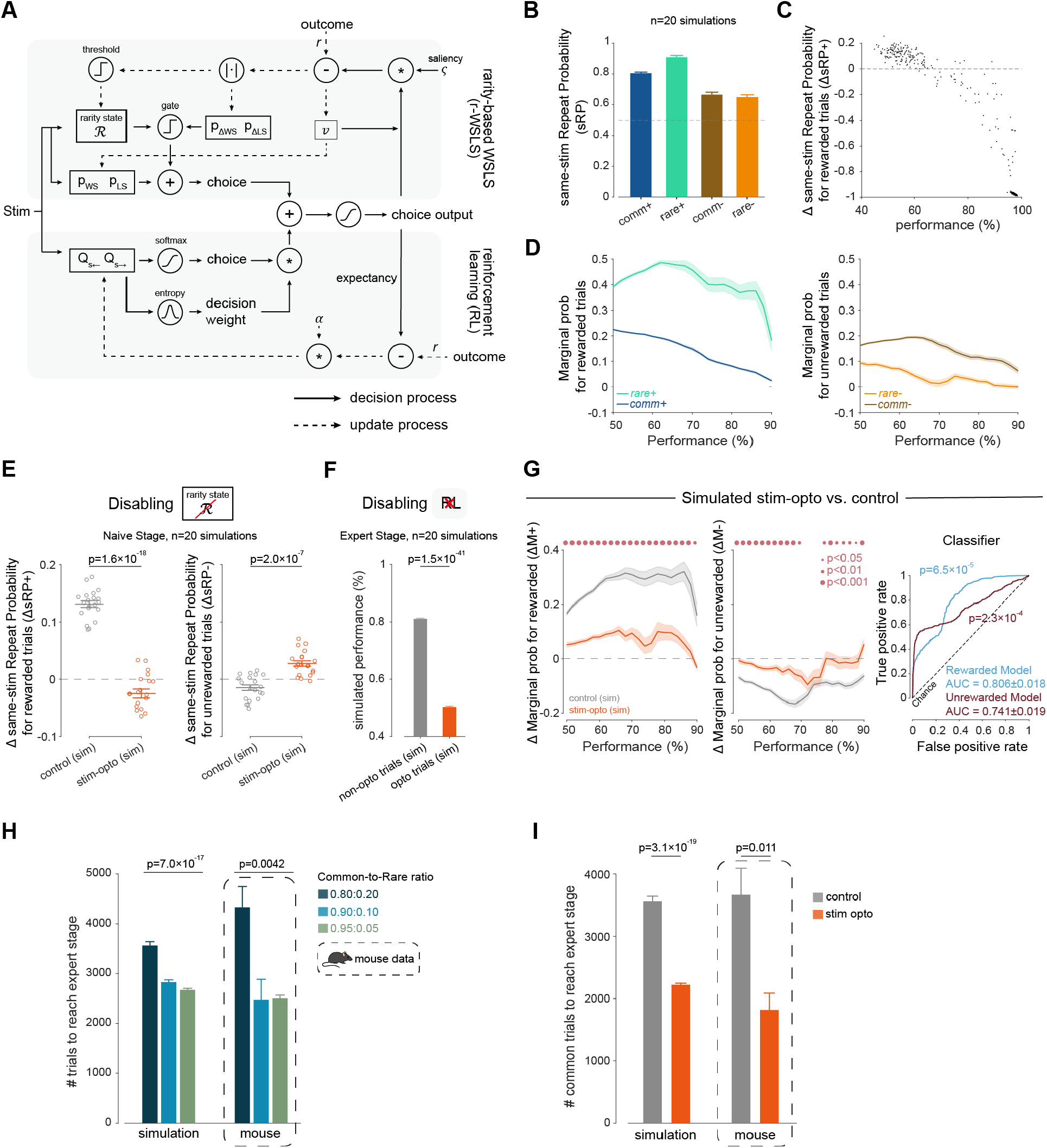
A data-driven metacognitive model recapitulates the decision bias and reveals PPC’s causal role in decision-making and learning under uncertainty. **(A)** The model consists of a rarity-based WSLS agent and a RL agent. **(B)** Simulated sRP (20 iterations) for 0.8:0.2 common-to-rare ratio. **(B)**Simulated ΔsRP_+_ across performance. **(C)** Simulated marginal probabilities for rewarded (left) and unrewarded (right) trials. **(E)** Simulated ΔsRP_+_ (left) and ΔsRP_-_ (right) by disabling rarity state in the WSLS agent in the naïve stage. **(F)** Simulated performance by disabling RL agent at the expert stage. **(G)** Simulated ΔM_+_ (left) and ΔM_-_ (middle) by disabling PPC functions in both WSLS and RL agents across learning. Right, ROC curves for classifiers predicting simulated *stim-opto* vs. control for rewarded and unrewarded trials. **(H)** Trial numbers to reach expert stage for 3 common-to-rare ratios, for model simulations (left) and mice (right). **(I)** Trial numbers to reach expert stage (common:rare=0.8:0.2) with PPC inactivation in rare trials, for model simulations (left) and mice (right). Wilcoxon ranksum test. Error bar, s.e.m.

This model accurately captured several key experimental findings. First, the model successfully recapitulated RI observed at 0.8:0.2, 0.9:0.1 and 0.95:0.05 common-to-rare ratios (**Figure 6B, Figure S10D**), as well as ΔsRP and ΔM (**Figure 6C, D, Figure S10D**). Second, disabling model components representing PPC functions reproduced findings from our optogenetics experiments: disabling the rarity state in the r-WSLS agent reduced ΔsRP_+_ to below 0 and increased ΔsRP_-_ to above 0 in the naïve stage (**Figure 6E**), disabling the RL agent impaired performance in the expert stage (**Figure 6F**), and disabling PPC functions in both agents reduced RI throughout learning (**Figure 6G, Figure S10E**). Moreover, the model predicted faster learning at lower rare ratio, closely resembling the learning speed observed in mice at 0.2, 0.1, and 0.05 rare ratios (**Figure 6H**). Finally, the model predicted that inactivating PPC in rare trials would accelerate learning, which was confirmed in our optogenetics experiments (**Figure 6I**), supporting a causal role of PPC in learning under uncertainty.

## Discussion

To navigate a world where uncertainty is pervasive, one needs to make good decisions in non-deterministic conditions. Using a probabilistic behavioral task with rare and common outcomes, we show that in both humans and mice, rare events exert a stronger impact on future decisions than common events, regardless of reward presence. Combining optogenetics and *in vivo* recordings, we demonstrate that PPC plays two distinct and opposing roles in this task: mediating the rarity-induced decision bias, which drives suboptimal choices, and driving stimulus-based decisions, which promotes optimal choices. These two functional roles involve overlapping PPC neuronal populations (**Figure 7**). Inspired by these results, we developed a computational model that captured PPC’s multiple functions in decision-making and learning under uncertainty.

**Figure 7.**
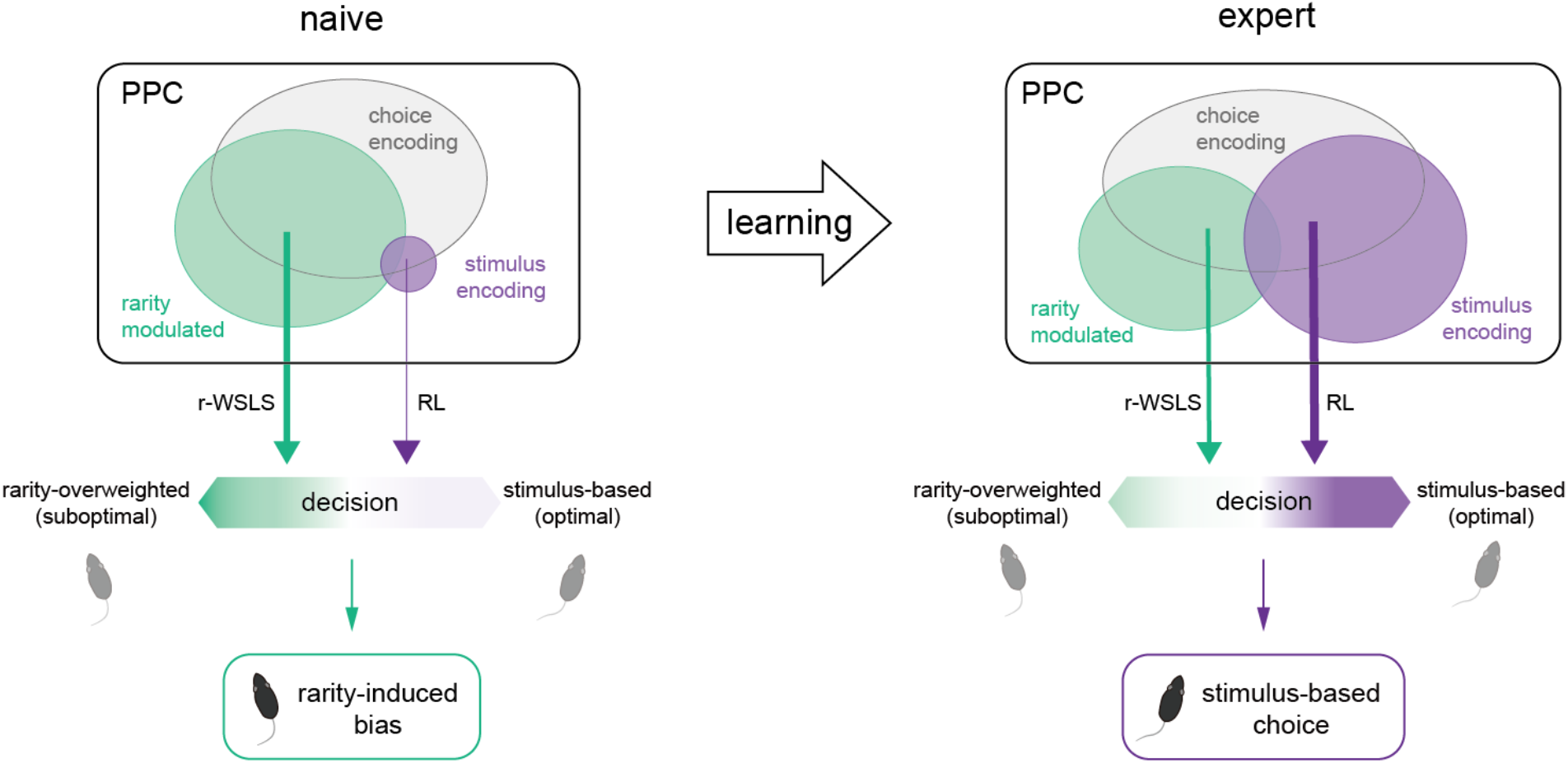
A diagram summarizing PPC’s dual functions in stimulus-based decision-making involving rare probabilities, during learning. Circles represent PPC neuronal populations that are rarity-modulated (green), stimulus-encoding (purple), and choice-encoding (grey), at naïve (left) and expert (right) stages. Animal’s choice for each trial is determined by two decision-making strategies, r-WSLS and RL, each corresponding to a PPC neuronal population. As learning progresses, the decision weight of r-WSLS decreases while that of RL increases. At the naïve stage, decision-making is strongly influenced by the suboptimal rarity-induced bias, whereas at the expert stage, decision-making is primarily stimulus-based, leading to optimal decisions.

Previous studies have established that judgments of rare probabilities tend to be systematically biased, influenced by features such as the recency, representativeness, saliency, and extremity of rare events^5,9-11^, and characteristics of decision-makers such as risk-seeking tendency, experience, and memory^5,10-13,16^. Risk and ambiguity of task parameters also influence decision-making involving rare probabilities^1,12,15^. By designing a task in which rarity is isolated from other factors, we identified and characterized a decision heuristic driven by probabilistic rarity itself, advancing our understanding of the behavioral phenomenon of overweighting rare probabilities. We found that this rarity impact is evident not only in the naïve stage, when subjects follow a stimulus-based WSLS strategy and perform the task at low accuracy, but also during the expert stage, when subjects have learned to make optimal decisions based on acquired stimulus-choice-reward associations. This finding highlights the robustness and persistence of the rarity impact in decision-making, echoing the idea that “the reliance on heuristics and the prevalence of biases are not restricted to laymen”^5^. This seemingly counterproductive bias may reflect an intrinsic, evolutionarily conserved preference for exploration over exploitation^8^, motivating animals to seek opportunities for new resources.

It was speculated that animals adopt different learning strategies for expected vs. unexpected uncertainty: fast learning under unexpected uncertainty (novel or changing environments), and slow learning under expected uncertainty (stable probabilistic contexts)^3,49^. Our data-driven model integrated both learning strategies: 1) in the early phase of the task, as subjects were not informed of the probabilistic nature of the task *a priori*, uncertainty was unexpected, and thus the fast-updating WSLS agent represented the fast learning strategy in novel environments; 2) as learning progressed and subjects became familiar with the task parameters, uncertainty became expected, and the slow-updating RL agent represented the slow stimulus-based associative learning in a stable probabilistic environment. To integrate these two agents, we used a metacognitive framework. Specifically, each agent’s contribution to the decision is weighted by its decision confidence, and the model transitions from WSLS-dominated in the naïve stage to RL-dominated in the expert stage. This model accurately reproduced the decision-making and learning behaviors, providing evidence for both cognitive strategies of learning under uncertainty. Furthermore, the metacognitive design made the model suitable for capturing complex learning processes in adaptive behaviors in various environments.

PPC is known to participate in perceptual decision-making in deterministic contexts^28,39^, and encode various task parameters in uncertain contexts^20,21,50^. However, evidence for PPC’s causal role in decision-making under uncertainty remains limited. Using optogenetics, we found two distinct roles that PPC played in decision-making under uncertainty: mediating the rarity impact, and driving stimulus-based decisions. *In vivo* recording of PPC neurons provided physiological evidence for both roles, by partially overlapping neuronal populations: rarity impact involves choice-encoding neurons modulated by rare events, some of which also encode auditory stimuli; stimulus-based decision-making involves stimulus-encoding neurons, many of which are also choice-encoding. In addition, the number of PPC neurons engaging in the task and their stimulus-encoding capacity evolve dynamically with learning, demonstrating PPC’s contribution to adaptive decision-making via learning-induced plasticity.

Our findings demonstrate that PPC encodes a higher-order history signal derived from recent experience, by integrating sensory, choice, and outcome information. Importantly, no single feature of rare trials (*i*.*e*., stimulus, choice, or outcome) in our study is rare on its own; rather, what is rare is the specific combination of these features. Our findings build on previous results that PPC encodes the history information of one or multiple task parameters^23-26^, and pinpoint PPC as a critical node for bridging complex past experience and future behavior, across different decision-making contexts. Future work is needed to establish a unified framework for PPC’s broad role in experience-dependent adaptive behavior.

## Resource availability

### Lead contact

Requests for further information and resources should be directed to and will be fulfilled by the lead contact, Yang Yang.

### Materials availability

This study did not generate new unique reagents.

### Data and code availability

All code is available at https://github.com/YangYangLab. All data and materials are available upon request.

## Acknowledgments

We thank Drs. Mu-ming Poo, Anthony M. Zador, Ju Lu, Qiaojie Xiong, and Danqian Liu for comments on the manuscript, and Drs. Balint Kiraly, Nicola Solari and Balazs Hangya for technical assistance on tetrode recording. We thank the animal facility, the imaging facility, and the HPC Platform of ShanghaiTech University for technical support. This work was supported by grants from the Ministry of Science and Technology of China (2022ZD0204900), Local Science and Technology Development Fund YDZX20233100001002, and ShanghaiTech University Startup Fund to YY, National Key R&D Program of China (2023YFA1009402) and the CAS Project for Young Scientists in Basic Research (YSBR-034) to MCT.

## Author contributions

Conceptualization: WS, YY. Methodology: WS, XH, YX, MCT, YY. Funding acquisition: MCT, YY. Visualization: WS, YY. Writing: WS, MCT, YY.

## Declaration of generative AI and AI-assisted technologies in the writing process

During the preparation of this work, the authors used Gemini (Google AI studio) in order to improve the language of the manuscript. After using this tool, the authors reviewed and edited the content as needed and take full responsibility for the content of the published article.

## Supplemental information

Figures S1–S10.

