## Supplemental figures for "Posterior parietal cortex mediates rarity-induced decision bias and learning under uncertainty"

**Figure S1, related to Figure 1**

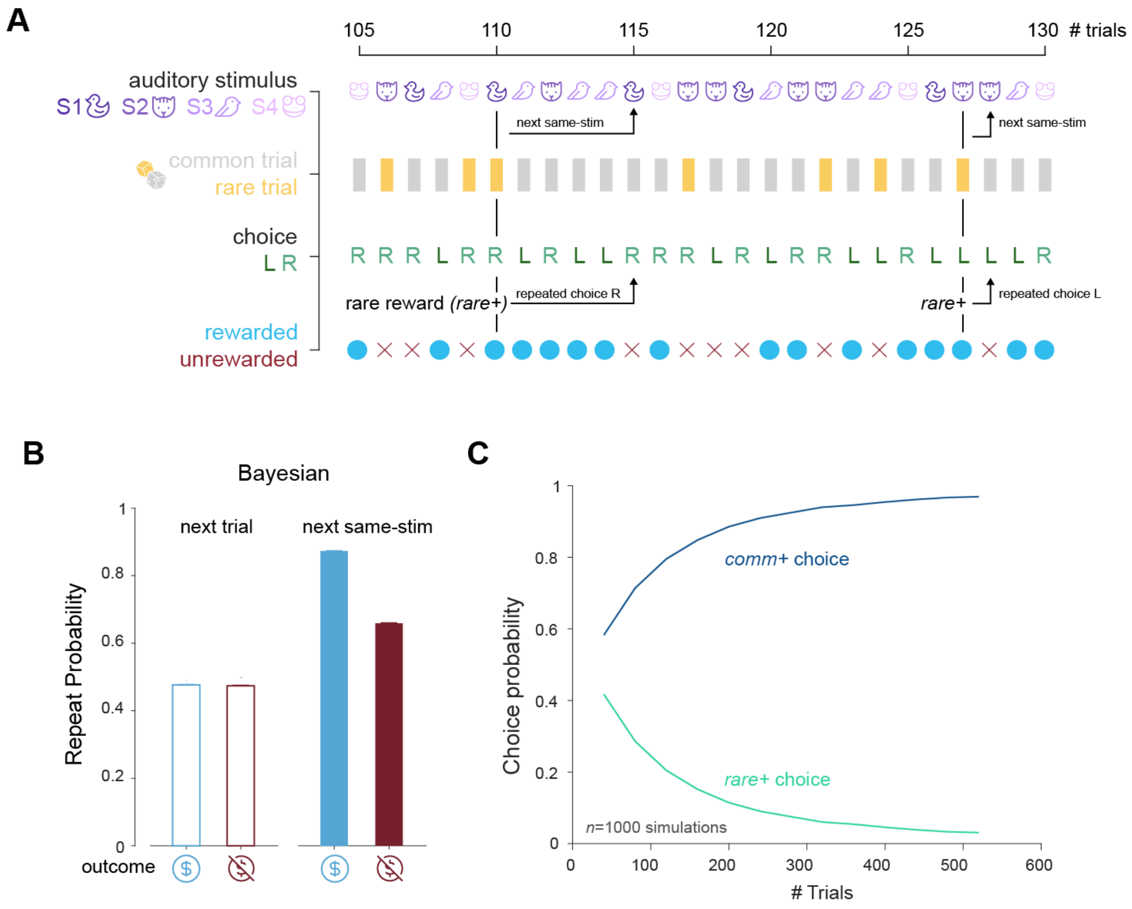


**Figure S1. An exemplary trial sequence of the task and Bayesian modeling results.**

(**A**) A trial sequence demonstrating the stimuli, the subject's choices, the outcomes, and the trial types (comm trial or rare trial) in the human behavioral task.

(**B**) Simulated choice repeating probabilities for the next trial (nRP) and the next same-stimulus trial (sRP), when the current trial is rewarded (blue) or unrewarded (red), of the Bayesian model.

(**C**) Simulated *comm*+ (correct) and *rare*+ (incorrect) choice probabilities of the Bayesian model.

#### Figure S2, related to Figure 2

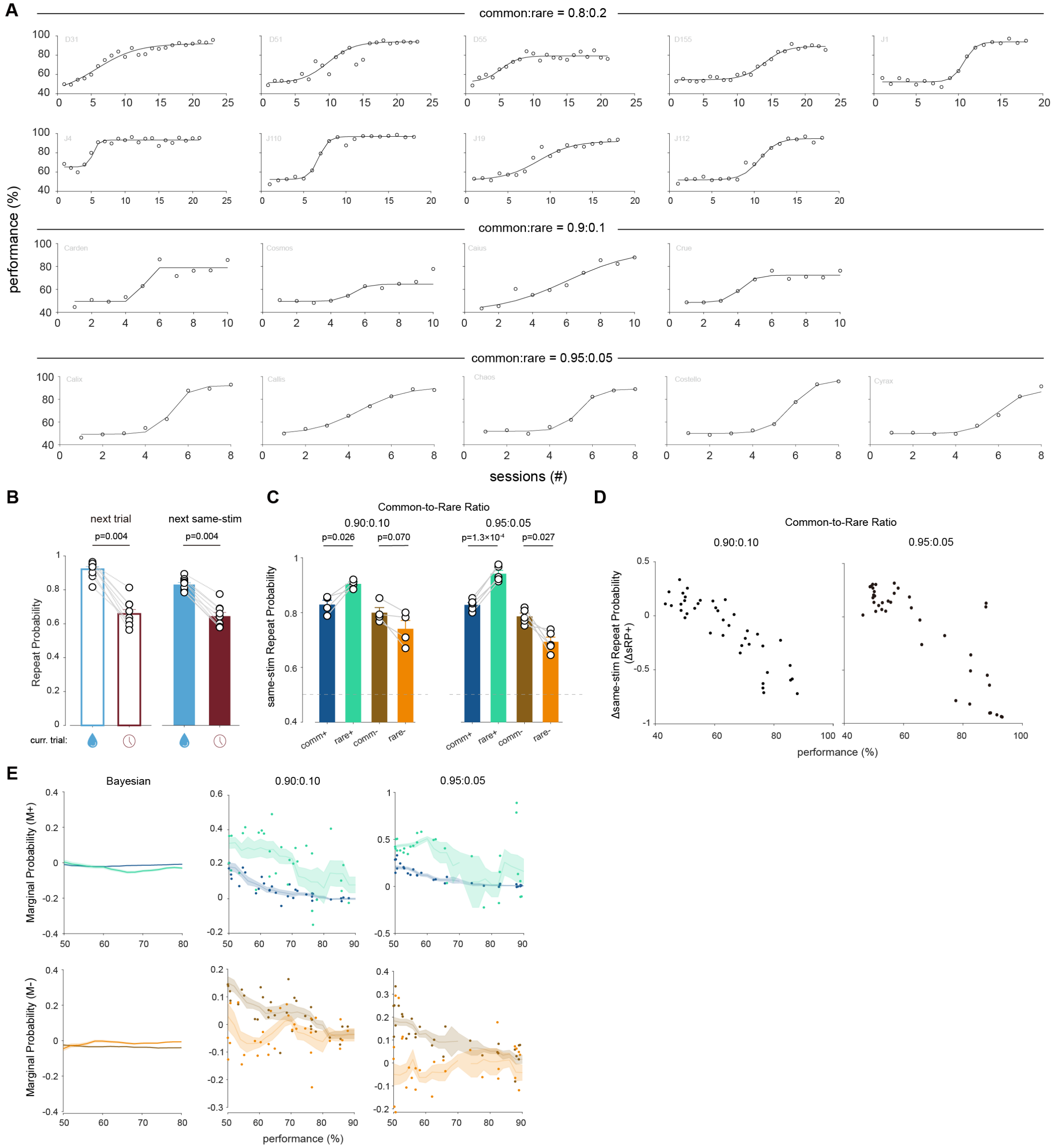


#### Figure S2. Mouse behavior results.

(**A**) Learning curves of all mice in 3 different common:rare groups, fitted to sigmoidal curves.

(**B**) Choice repeating probability for the next trial and next same-stimulus trial, when the current trial is rewarded (blue) or unrewarded (red), in the 0.8:0.2 group. Wilcoxon signed-rank test.

(**C**) sRP for 0.9:0.1, and 0.95:0.05 mice in the naïve stage.

(**D**) $\Delta sRP_+$ across session for 0.9:0.1, and 0.95:0.05 mice.

(**E**) $M_{rare+}$ and $M_{comm+}$ (top), $M_{rare-}$ and $M_{comm-}$ (bottom) for Bayesian model (left), 0.9:0.1 mice (middle) and 0.95:0.05 mice (right).

**Figure S3, related to Figure 3**

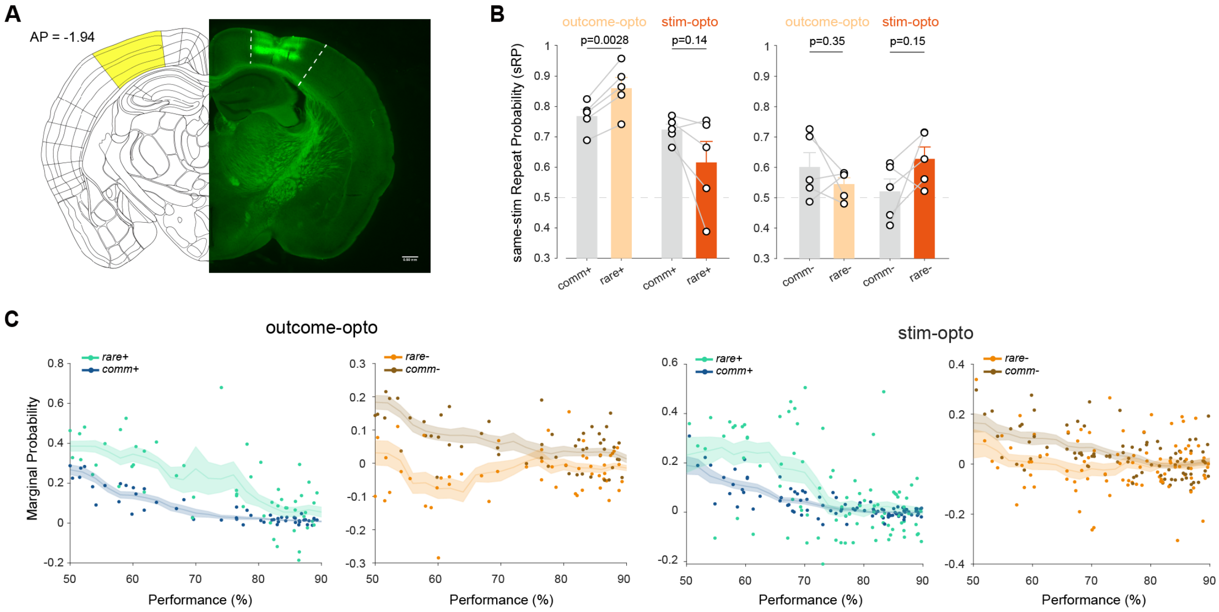


### Figure S3. Optogenetic inhibition of PPC.

(**A**) Histology showing the optogenetic inhibition site with side-by-side comparison with the mouse brain atlas (Allen Atlas). Scale bar, 0.5mm.

(**B**) Bar graph showing sRP for *outcome-opto* and *stim-opto* mice in the naïve stage. Each data point represents one mouse. Wilcoxon signed-rank test.

(**C**) Left, $M_{rare+}$ and $M_{comm+}$, $M_{rare-}$ and $M_{comm-}$ for *outcome-opto* mice. Right, $M_{rare+}$ and $M_{comm+}$, $M_{rare-}$ and $M_{comm-}$ for *stim-opto* mice.

**Figure S4, related to Figure 4**

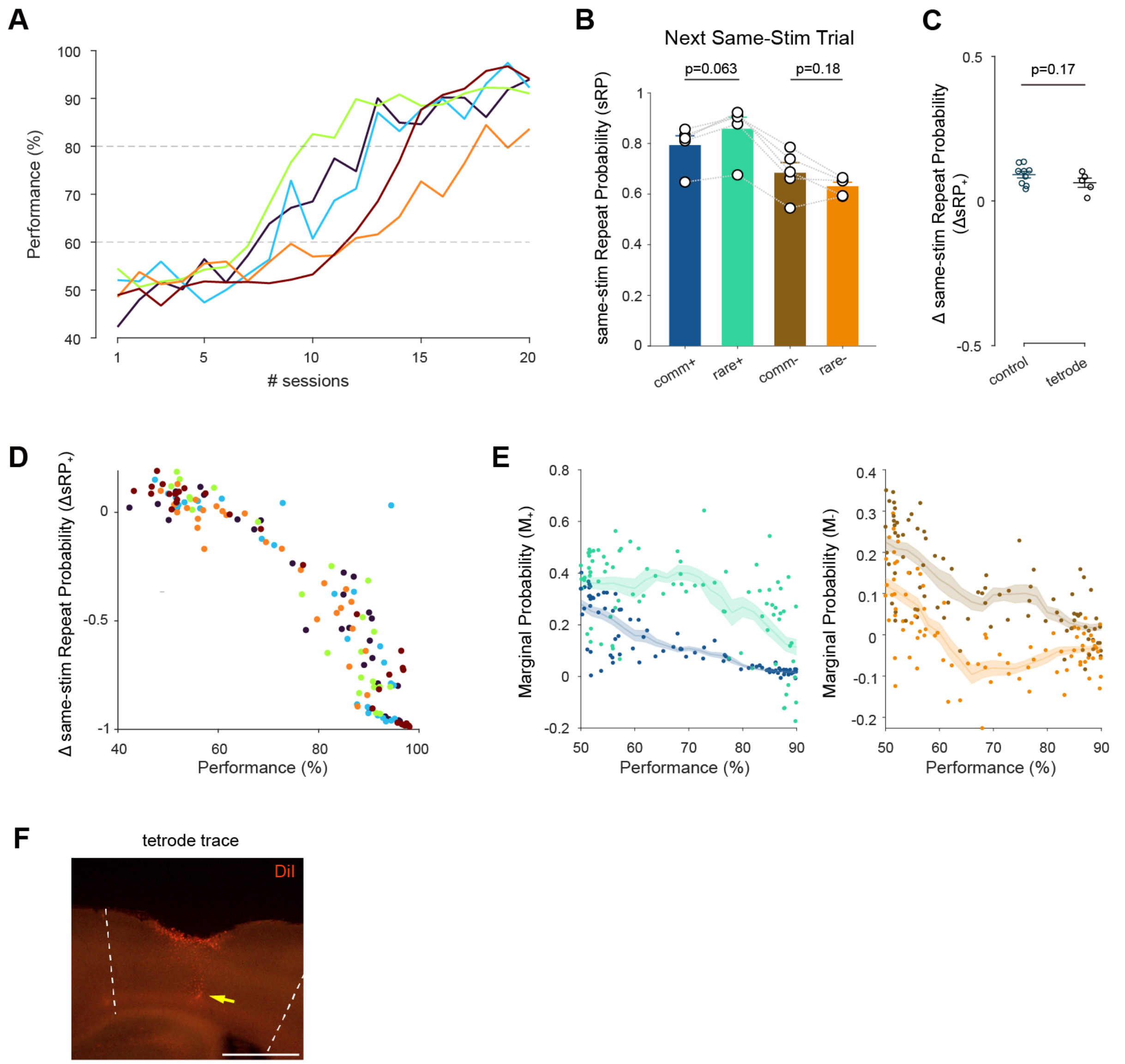


**Figure S4. Behavior results of mice used in tetrode recording experiments.**

(**A**) Performance of all 5 mice used in tetrode recording experiments across sessions. Each line represents one mouse.

(**B**) sRP for tetrode recording mice in the naïve stage. Each data point represents one mouse.

(**C**) ΔsRP of tetrode recording mice in the naïve stage is similar to ΔsRP of control mice.

(**D**) ΔsRP across performance for tetrode recording mice.

(**E**) $M_{rare+}$ and $M_{comm+}$ (left), $M_{rare-}$ and $M_{comm-}$ (right) for tetrode recording mice.

(**F**) Histology showing the tetrode trace in the PPC. Tetrodes were dipped in DiI before implantation. Yellow arrow points to the deepest recording location. Scale bar, 1 mm.

#### Figure S5, related to Figure 4

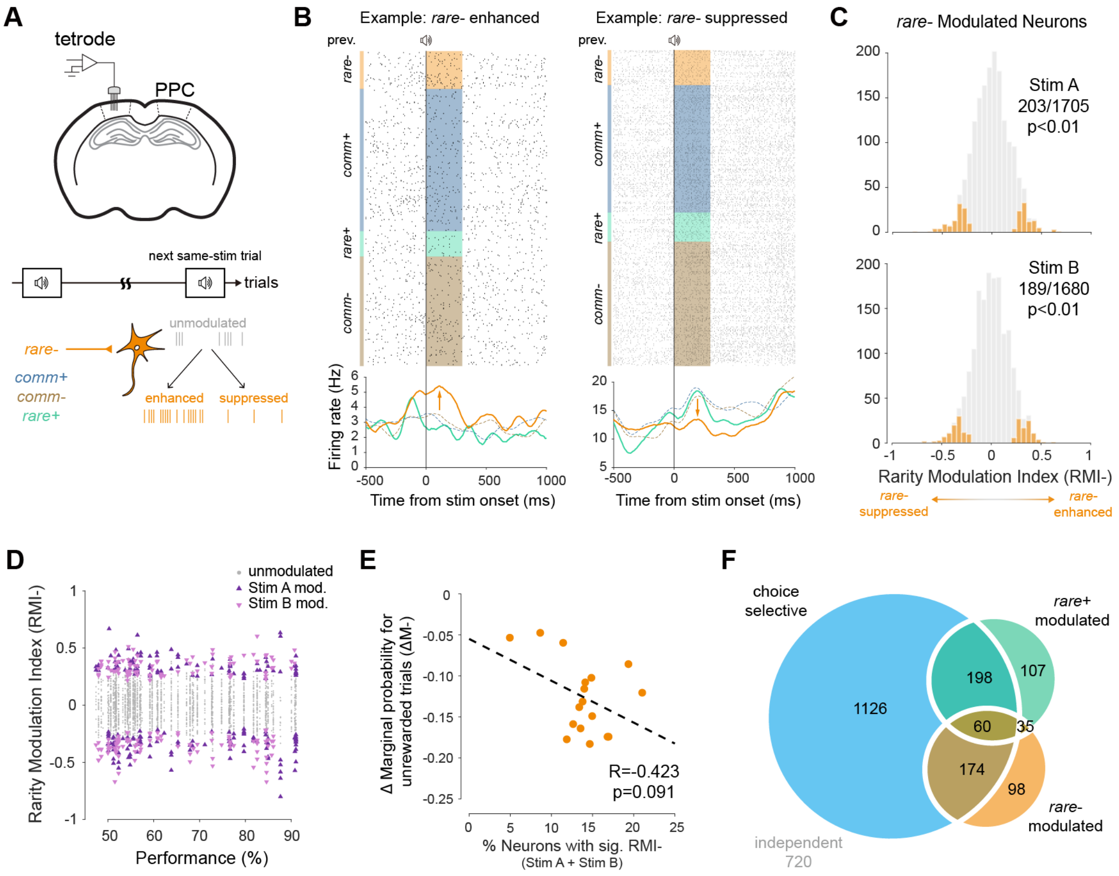


**Figure S5. *Rare*- modulation of PPC neurons.**

(**A**) Top panel, schematic showing tetrode recording in PPC. Bottom panel, schematic showing modulation by preceding *rare*- trials on PPC neuronal activity.

(**B**) Spike raster and peri-stimulus time histogram (PSTH) of 2 example neurons, either enhanced (left) or suppressed (right) by preceding *rare*-.

(**C**) Histogram showing *rare*- modulation index ($RMI_-$) of PPC neurons calculated using ROC analysis, for stimulus A (top panel) and stimulus B (bottom panel). Color bars represent significant modulation (bootstrapping, 200 iterations).

(**D**) $RMI_-$ of individual neurons recorded across performance. Triangles, neurons significantly modulated by stimulus A or stimulus B *rare*-.

(**E**) Correlation between the RI strength for *rare*- trials ($\Delta M_-$) and the fraction of neurons with significant $RMI_-$.

(**F**) Overlap of choice-selective, *rare*+ modulated and *rare*- modulated PPC neurons.

**Figure S6, related to Figure 4**

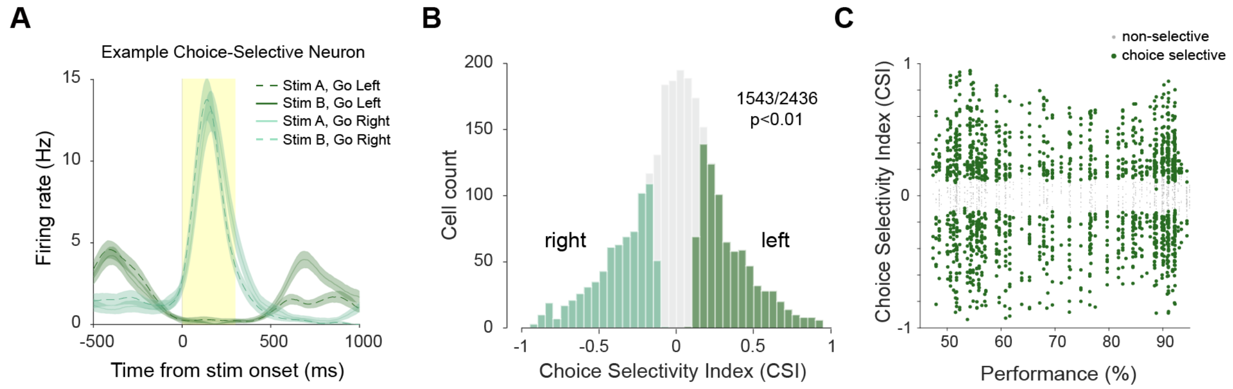


**Figure S6. Choice selectivity of PPC neurons.**
(**A**) Peri-stimulus time histogram (PSTH) of an example right-selective PPC neuron.
(**B**) Histogram showing choice selectivity index (CSI) of PPC neurons. CSI was calculated using ROC analysis. Colored bars represent significant choice-selective neurons. Statistical significance was assessed via bootstrapping (200 iterations, $p<0.01$).
(**C**) CSI of individual neurons recorded across performance. Green dots, neurons significantly selective to either left (CSI>0) or right (CSI<0).

#### Figure S7, related to Figure 4

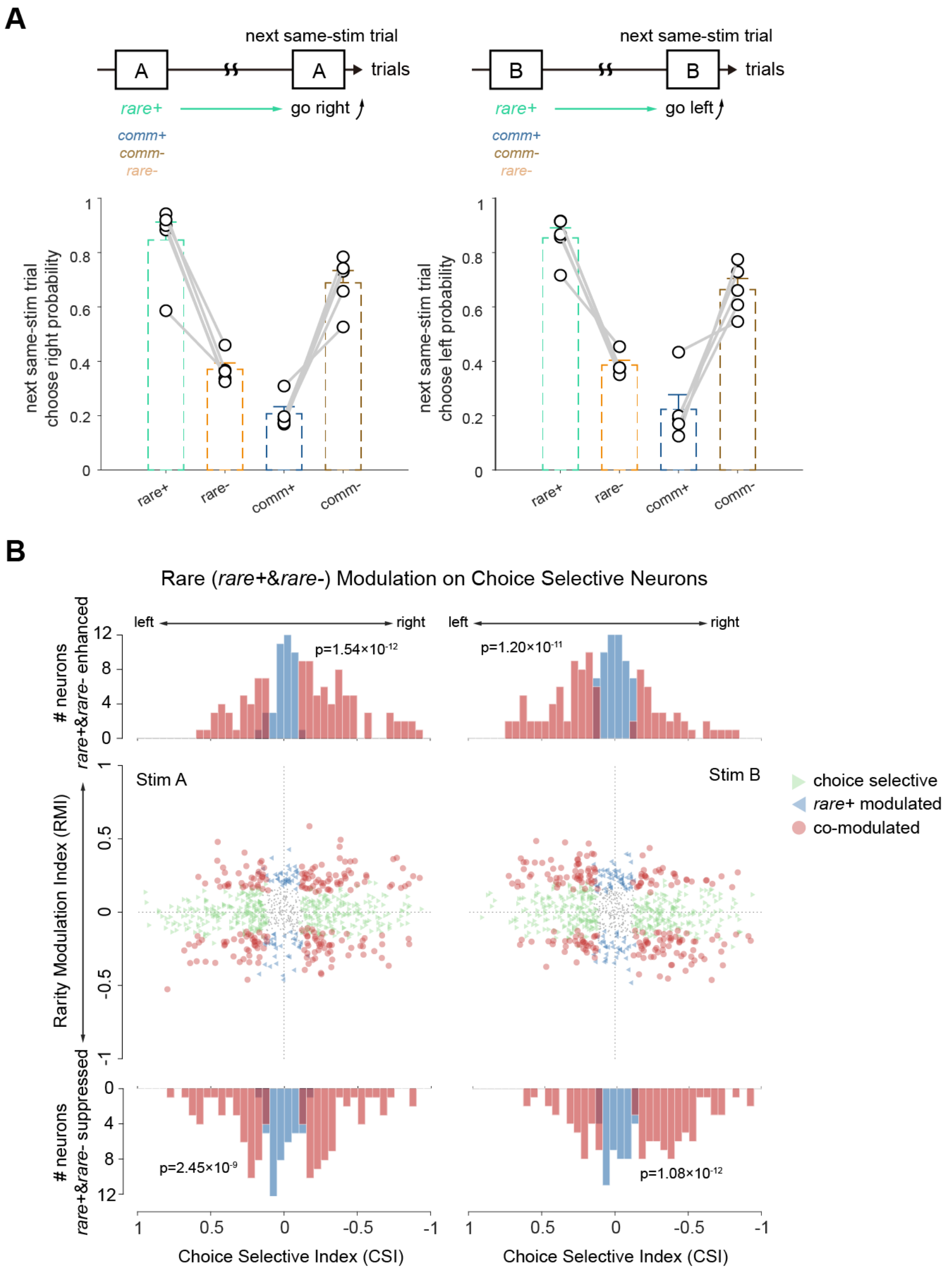


**Figure S7. Rarity modulation of PPC neurons.**

(**A**) Left panel, stimulus A *rare+* (right-going *rare+* trial) yields the highest probability of going right in the next stimulus A trials. Right panel, stimulus B *rare+* (left-going *rare+* trial) yields the highest left-going probability in the next stimulus B trials.

(**B**) RMI (including both *rare+* and *rare-*) plotted against CSI for all neurons. Left panel, stim A; right panel, stim B. Each data point represents one neuron. Green triangle, choice-selective neurons. Blue triangle, rarity modulated neurons. Red dots, choice and rarity co-modulated neurons. Histograms show CSI for rarity enhanced (top) and rarity suppressed (bottom) neurons, respectively. Kolmogorov-Smirnov test.

**Figure S8, related to Figure 5**

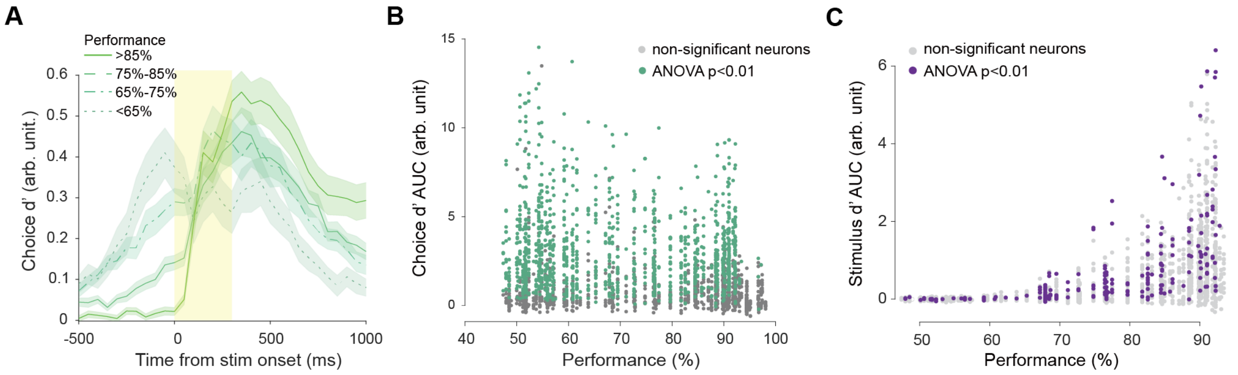


**Figure S8. Choice and stimulus discriminability (d') of PPC neurons.**
(**A**) Averaged choice d' for significant choice-encoding neurons recorded in sessions at different performance levels.
(**B**) Area under curve (AUC) of choice d' for choice-encoding neurons across performance levels.
(**C**) AUC of stimulus d' for stimulus-encoding neurons across performance levels.

#### Figure S9, related to Figure 6

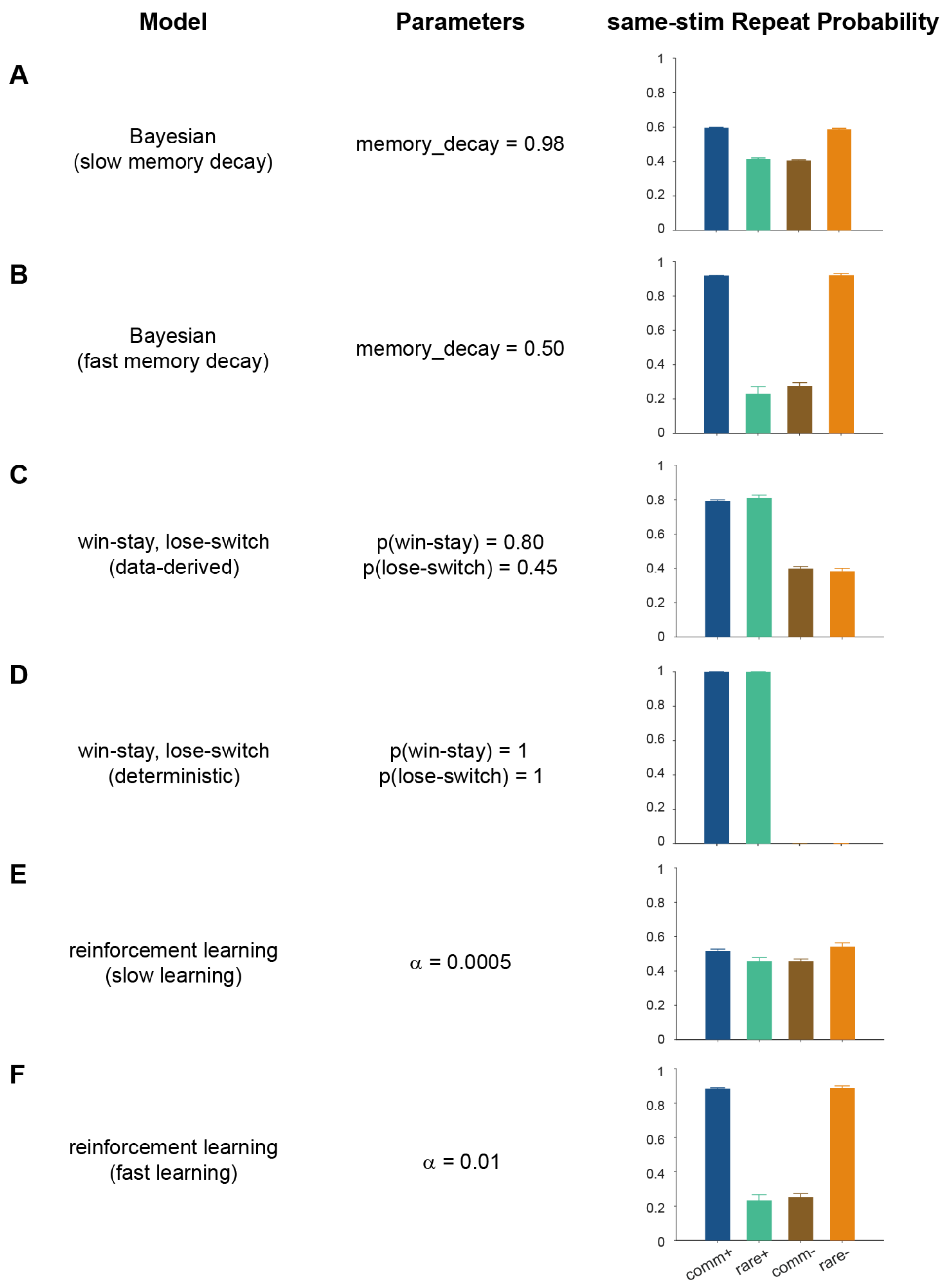


**Figure S9. Simulation results for Bayesian, WSLS, and RL models.**
(**A**) Simulated sRP for a Bayesian model with slow memory decay.
(**B**) Simulated sRP for a Bayesian model with fast memory decay.
(**C**) Simulated sRP for a WSLS model with parameters derived from behavioral data.
(**D**) Simulated sRP for a WSLS model in which the agent always stays after a win and switches after a loss.
(**E**) Simulated sRP for a RL model with slow learning rates.
(**F**) Simulated sRP for a RL model with fast learning rates.

#### Figure S10, related to Figure 6

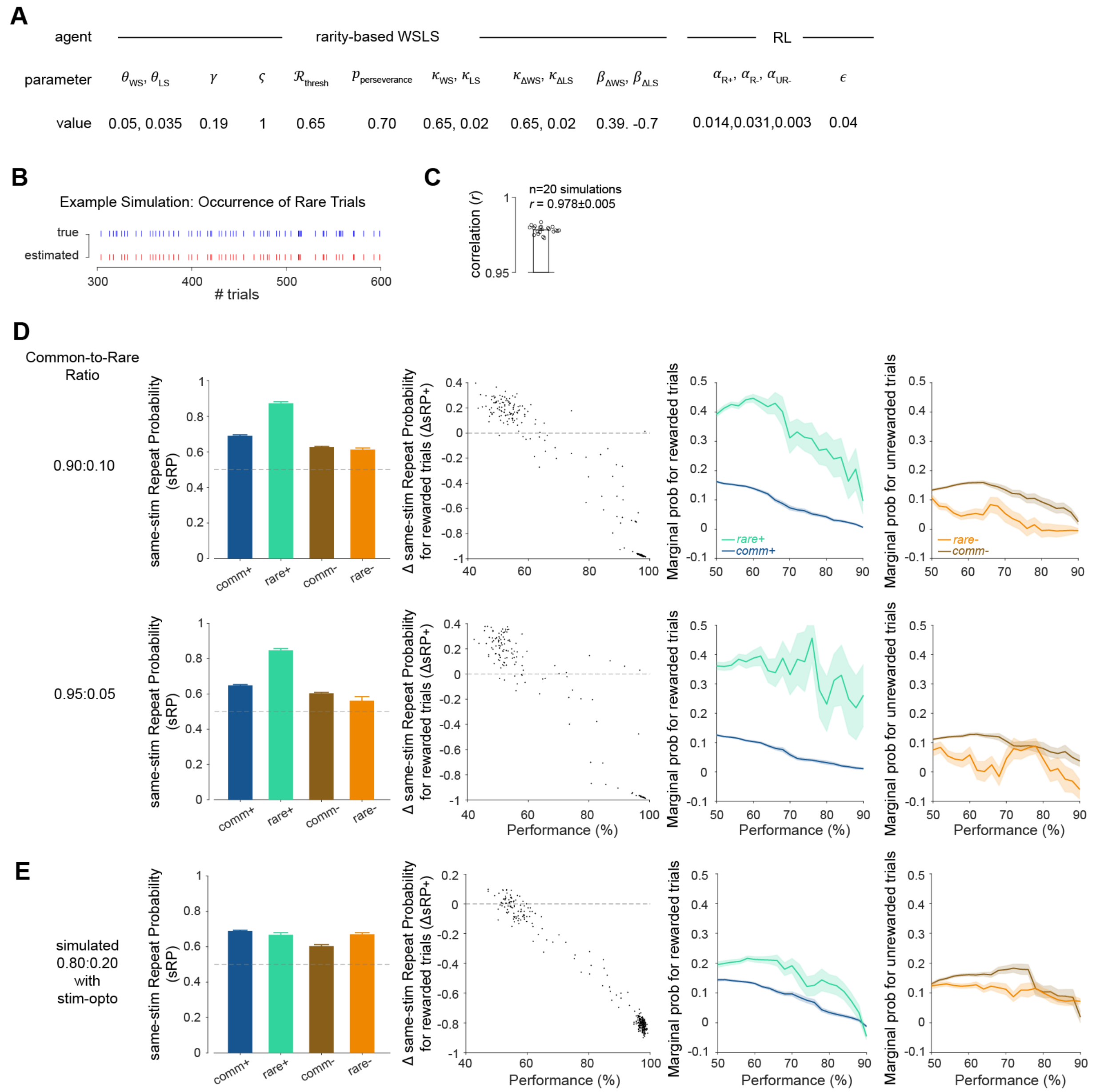


**Figure S10. Metacognitive modeling results.**

(**A**) Parameters and values used in the model.

(**B**) Example trials showing the actual occurrence of rare trials and estimated occurrence by the model, given the trial sequence.

(**C**) Correlation of actual and estimated rare trial occurrences.

(**D**) Simulated sRP, $\Delta$sRP$_+$, $M_{rare+}$ and $M_{comm+}$, $M_{rare-}$ and $M_{comm-}$, $\Delta M_+$ and $\Delta M_-$ for 0.9:0.1, and 0.95:0.05 common-to-rare ratios.

(**E**) Simulated sRP, $\Delta$sRP$_+$, $M_{rare+}$ and $M_{comm+}$, $M_{rare-}$ and $M_{comm}$, for simulated inactivation of PPC, by disabling PPC functions in both model agents.
