## Supplementary material for "Posterior parietal cortex mediates rarity-induced decision bias and learning under uncertainty": Methods

#### Human subjects

A total of 13 human subjects (4 males and 9 females; mean age ± s.d.: 24.0±5.2, range: 19–40) were recruited in the study. All subjects were healthy and had normal vision and hearing. This study was approved by the Research Ethics Committee of ShanghaiTech University. Every subject signed an informed consent and received monetary compensation for participation. Subjects were informed that the study was related to learning and memory in general, and were instructed to make auditory stimulus-guided decisions for monetary rewards. Subjects were not aware of the probabilistic task design prior to participation.

#### Human behavioral task

Subjects were seated in front of a computer with a speaker, an LCD monitor, and a custom-made keypad with three keys: "S" for start, "L" for the left choice, and "R" for the right choice. "S" key is controlled by the subject's left hand, and the two choice keys ("L" and "R") are controlled by the subject's right hand.

Before the task, subjects were allowed to adjust the monitor, speaker and keypad to comfortable positions. Auditory stimuli were played from the speaker, controlled using the PsychToolbox software[1]. Key presses were recorded using the Bpod system (Bpod State Machine r2.5, Sanworks) using custom-made software (Matlab). Task instructions were given as texts on the monitor screen. Subjects first went through a pre-task stage. In this stage, auditory stimuli (pure tones, not used in the real task) were played, and subjects were instructed to press the L or R key as specified on the screen upon hearing the auditory stimuli.

Subsequently in the task, subjects were instructed to obtain monetary rewards by pressing the L or R key based on auditory stimuli. Four distinct auditory stimuli are used: duck, cat, canary, and frog sounds (mp3 files downloaded from pixabay.org). The subject presses the "S" key to initiate a trial, triggering the delivery of 1 of 4 auditory stimuli. Upon hearing the stimulus, the subject makes a choice by pressing the "L" or "R" key. As soon as the choice is made, a cartoon showing money earned (rewarded) or money lost (unrewarded) is displayed on the monitor screen. For a duck or cat sound, "L" choice is rewarded at 0.8 probability (common), and "R" choice is rewarded at 0.2 probability (rare); For canary or frog sound, "R" choice is rewarded at 0.8 probability (common), and "L" choice is rewarded at 0.2 probability (rare). Inter-trial interval lasts 2-6 seconds. Stimuli and common/rare trials are pseudorandomly interleaved: the same stimulus will not be repeated consecutively for more than 5 trials; a rare trial is always followed by a common trial.

#### Human behavioral analysis

We defined choices with the higher reward probability as correct choices for calculating task performance: "L" for "duck" and "cat" sounds, and "R" for "canary" and "frog" sounds are considered correct. Performance is calculated as the percentage of correct choices in 20 consecutive trials.

Ten out of 13 subjects were able to achieve >85% performance after 60~400 trials. The task is terminated when subjects reached 85% performance. To define the naïve stage, we used an empirical performance threshold of 70%. At least 6 *rare+* trials were included in the analysis for each subject. Two subjects (1 male and 1 female) encountered only 2 and 3 rare+ trials in the whole session, respectively, and were excluded from the analysis.

The same-stim Repeat Probability (sRP) was defined as the probability of repeating the same choice in the next same-stimulus trial, and is further divided into four variables based on the four possible outcomes: *comm+*, *rare+*, *comm-*, and *rare-*. ΔsRP is then consecutively defined as the differences in sRP following rewarded trials and unrewarded trials, and is defined as:

$$\Delta sRP_{+} = sRP_{rare+} - sRP_{comm+}, \text{ for rewarded trials,}$$

$$\Delta sRP_{-} = sRP_{rare-} - sRP_{comm-}, \text{ for unrewarded trials.}$$

To compute the performance, ΔsRP, and ΔM, trials were first divided into 40-trial blocks for each subject. Note that these were all calculated for the next same-stimulus trials, not the next trials.

To calculate $M_{rare+}$ and $M_{comm+}$:

$$M_{rare+} = P(\text{incorrect choice} \mid \text{preceding } rare+) - P(\text{incorrect choice})$$

$$M_{comm+} = P(\text{correct choice} \mid \text{preceding } comm+) - P(\text{correct choice})$$

Here, P(incorrect choice|preceding *rare+*) is the conditional probability of making *rare+* choice after a *rare+* trial (i.e., $sRP_{rare+}$), P(incorrect choice) is the probability of making *rare+* choice after any trial in the same session. P(incorrect choice) = 1-performance. Similarly, P(correct choice |preceding *comm+*) is the conditional probability of making *comm+* choice after a *comm+* trial (i.e., $sRP_{comm+}$), P(correct choice) is the probability of making *comm+* choice after any trial: P(correct choice) = performance. This gives:

$$M_{rare+} = sRP_{rare+} - (1 - \text{performance})$$

$$M_{comm+} = sRP_{comm+} - \text{performance}$$

To compare $M_{comm+}$ and $M_{rare+}$, smoothed $M_{comm+}$ and $M_{rare+}$ were computed by averaging $M_{comm+}$ and $M_{rare+}$ around each performance level with a ±5% window, respectively.

### Animals

Wild-type C57BL/6 mice were purchased from Shanghai Jihui Laboratory Animal Care Co., Ltd. Mice were housed and bred in a 12 h light-dark cycle (7 am - 7 pm light) in the animal facility of ShanghaiTech University. Eight- to 16-week-old male mice were used in the experiments. Animal protocols were approved by the Research Ethics Committee of ShanghaiTech University.

### Mouse behavioral task

All behavioral experiments were conducted in a custom-made, electromagnetic shielded, sound-proof booth[2]. A custom-made behavioral chamber has three ports (Mouse port assembly, Sanworks) in the front. Eighteen mice were trained in the task. Mice had limited water access each day starting from three days prior to behavioral training, and were weighed every day to maintain at least 80% of the original body weight. A simple pre-training task was introduced to familiarize the mice with the setup. Specifically, mice were primed with a neutral sound (8 kHz pure tone) to learn the poking sequence: poking into the center port triggers an auditory stimulus, and poking into either side port (left or right) always yields a water reward. This pre-training takes 2 sessions. In the 2AFC task with probabilistic outcomes, the mouse first pokes into the center port to trigger the delivery of an auditory stimulus (stimulus A: 5 kHz pure tone, 65dB; stimulus B: 13 kHz pure tone, 65dB). The stimulus is preceded by a random (uniform distribution) delay of 50-100ms to prevent the mice from making preliminary decisions, and is terminated when the mouse pokes into a side port (left or right). The outcome (3 μL water reward or 3-second time-out) is delivered immediately as soon as the mouse pokes into a side port. Stimulus A is associated with reward in the left port with 0.8 probability (common) and the right port with 0.2 probability (rare); stimulus B is associated with reward in the right port with 0.8 probability (common) and the left port with 0.2 probability (rare). Inter-trial interval lasts 1-2 seconds. Stimuli and common/rare trials are pseudorandomly interleaved: the same stimulus will not be repeated consecutively for more than 5 trials; a rare trial is always followed by a common trial.

**Mouse behavioral analysis**

We defined choices with the higher reward probability as correct choices for calculating task performance: left choice for stimulus A and right choice for stimulus B are considered correct. Performance was calculated as the percentage of correct trials in each behavioral session. Each session typically contains 300~600 trials. $M_{rare+}$ and $M_{comm+}$ are computed the same way as in humans, except that all calculations were based on sessions, rather than 40-trial blocks. Smoothed M, ΔsRP and ΔM were computed by averaging ΔsRP and ΔM around each performance level with a ±5% window, respectively.

**Classifier of behavioral groups**

We built a classifier using MATLAB's fitcdiscr function (quadratic discriminant analysis, 5-fold cross-validation) to discriminate between groups of behaviors [3]. The average receiver operating curves (ROCs) were shown to demonstrate each classifier's performance. The area under curve (AUC) was calculated using MATLAB's perfcurve, and the 5-fold AUC values were compared to 0.5 (chance) to produce p-value using t-test.

**Bayesian modeling**

To model the decision-making and learning process of humans and mice in the probabilistic task without the rarity impact, we implemented a simple Bayesian reinforcement learning model. This model maintains a belief about the probability of reward associated with choosing the left and right options for a given stimulus, represented by beta distributions. The prior beliefs about the reward probabilities ($p_L$ and $p_R$) are parameterized by $\alpha$ and $\beta$ for each choice and stimulus type. On each trial, after the presentation of a stimulus, the model randomly samples from the beta distributions to estimate the expected value for each choice ($c_L$ and $c_R$). The model then generates a choice by combining these sampled values with added Gaussian noise, such that the left choice is made if $c_L - c_R + \epsilon > 0$, where $\epsilon \sim N(0, \sigma^2)$. After receiving the outcome, the priors $p_L$ and $p_R$ are first multiplied by a discounting factor to represent evidence decay, and then updated based on the Bayesian rule to produce the posterior beliefs, incorporating the observed outcome of the current trial. Briefly, a rewarded choice increases the $\alpha$ parameter, shifting the posterior distribution towards higher probabilities of reward for that choice, while an unrewarded choice increases the $\beta$ parameter, shifting the distribution towards lower probabilities of reward.

**Virus injection and fiber implantation**

For the optogenetic inhibition experiments, the 594nm-activated halorhodopsin (eNpHR3.0) fused with yellow fluorescent protein (YFP) is expressed in mouse PPC by stereotaxic injection of adeno-associated virus (AAV9-CaMKII-eNpHR3.0-eYFP, $4.5 \times 10^{13}$ v.g./mL, OBiO Technology). Mice were anesthetized with isoflurane (1%-1.5%) and positioned in a stereotaxic device (Reward

Co.). Body temperature was maintained at 37°C using a small heating pad. Virus was first diluted to $4.5\times10^{12}$ v.g./mL with cell-grade PBS (Solarbio), and injected using a glass pipette with a tip diameter of 20–25 μm via a small skull opening (<0.4 $mm^2$) with a micro-injector (Nanoject3, Drummond). An aliquot of 300 nL was injected into the PPC of both hemispheres at 1nL/s. The stereotaxic coordinates were: AP -1.94mm, ML ±1.60mm, DV 0.50mm. Optic fibers (0.37 N.A., 200 μm core diameter, 1mm length, Inper) were implanted at a depth of 200 μm below cortical surface at the viral injection sites 2 weeks after virus injection. Following the fiber implantation, mice recover for 1 week before the behavioral training started.

**Optogenetic experiments**

Ten mice were used for the optogenetic experiments (N=5 for PPC inhibition during the stimulus epoch, N=5 for PPC inhibition during the outcome epoch). Before each session, the implanted optic fiber was connected to a 200μm-core connected fiber optic (Inper) and a rotary joint (Inper) to allow free moving without restriction by the cable. A 589-nm DPSS laser (YL589T6, SLOC) was used, and the laser power was measured before each session to ensure a laser delivery of ~10mW at the fiber tips.

For optogenetic inhibition during the outcome epoch (*outcome-opto*): laser delivery was triggered by the side poke, and lasted until the mouse left the side port or a minimum of 3 seconds in unrewarded trials. To ensure that mice did not treat the laser illumination as an additional cue, a masking yellow light was delivered during the outcome epoch of all trials.

For optogenetic inhibition during the stimulus epoch (*stim-opto*): laser delivery was triggered by the center poke, and lasted until the mouse poked into one of the side ports, the same time as the termination of the auditory stimulus. A masking yellow light was delivered during the stimulus epoch of all trials.

**In vivo electrophysiology**

Custom-made tetrode drives were assembled using polyimide-coated nichrome tetrode wires (PX000004, Sandvik), a 32-channel EIB board (EIB-36-PTB, Neuralynx), and a 3-d printed skeleton that altogether weighed ~0.8 g. Eight tetrodes were gold-plated to impedance <200 kW prior to implantation, and implanted in the PPC of both hemispheres. The tetrode tips were dipped in DiI solutions (MB4240-1, meilunbio) to allow post-mortem checking of the implantation site. For tetrode drive implantation, mice were anesthetized with isoflurane (1%-1.5%) and positioned in a stereotaxic device (68803, Reward Co.). Body temperature was maintained at 37°C using a small heating pad. The tetrodes were implanted at the stereotaxic site of AP -1.94mm, ML ±1.60mm, DV 0.50mm. Mice recover in their home cages for 1 week before the training started.

Five mice were used for the tetrode recording experiments. Electrophysiological data was collected using the OpenEphys recording system (OEPS-9030, OpenEphys). Before each session, the tetrode drive was connected to a headstage (#C3314, Intan) and the OpenEphys system using a soft connector wire (#C3216, Intan). A custom-made rotator was used to allow free moving of the mice. Before each recording session, each tetrode was independently advanced by 10-40 μm to enable recording of different neurons. To detect spiking activity, the recorded data were bandpass filtered between 600 Hz and 6 kHz, and spikes were thresholded at 40 μV. Clusters were manually sorted using MClust 3.5 (https://redishlab.umn.edu/mclust) by expert experimenters to derive single units. A unit was deemed valid and included in the analysis if spikes had <1% refractory period violations.

### Electrophysiological data analysis

Single units with mean firing rates of < 2 Hz were excluded from all analyzes. PSTHs were calculated by averaging firing rates in 10-ms bins and smoothed with a Gaussian kernel ($\sigma = 50$ ms).

***Rare*+ modulation index**. To quantify how neural activity of each unit was affected by previous outcomes (*rare*+ vs. others), we used an ideal observer decoding based on receiver operating characteristics (ROC) analysis[4]. Using the activity of the stimulus epoch, we defined a *rare*+ modulation index (RMI) as 2 × (auROC – 0.5), which ranged from -1 to 1. $RMI_+ > 0$ indicates higher firing rates following *rare*+ trials than following other trials (*rare*+ enhanced); $RMI_+ < 0$ indicates lower firing rates following *rare*+ trials than following other trials (*rare*+ suppressed). A minimum of 3 *rare*+ trials are required for a cell to be included in the analysis. The significance was assessed via bootstrapping (200 iterations).

***Rare*- modulation index**. Same as *rare*+, except that $RMI_-$ was computed on firing rates following *rare*- trials vs. other trials.

**Choice selectivity index**. To quantify how each unit's activity encodes the current choice, we estimated auROC on the activity of the stimulus epoch, and defined a choice selectivity index (CSI) as 2 × (auROC – 0.5). CSI>0 indicates left selectivity, and CSI<0 indicates right selectivity. Significance was assessed via bootstrapping (200 iterations).

**ANOVA**. To quantify the significance of encoding stimuli and choices, we used two-way ANOVA to fit neural activity of the stimulus epoch to stimuli and choices. A neuron is considered significantly stimulus- or choice-encoding if the corresponding $p < 0.01$.

**Stimulus and choice discriminability (d')**. To compute stimulus d', we first computed the mean difference in the smoothed, 50-ms-binned firing rate on the two different trial types (stimulus A vs. B) divided by the square root of their mean variance [5]. Bootstrapping was performed after balancing trial numbers. We then subtracted this value with the mean shuffled d' (200 iterations). To compute choice d', we computed the mean difference in firing rate on the two different trial types (Left vs. Right) divided by the square root of their mean variance, then subtracted this value with the mean shuffled d' (200 iterations).

**Demixed principal component analysis.** We followed the procedure in [6], and used the code provided by the authors to analyze our data (https://github.com/machenslab/dPCA). The multi-dimensional input matrices are organized as: neuron ID * stimulus type (A or B) * choice type (L or R) * time range * trial ID. The time range used for analysis is 0~300 ms after stimulus onset. A minimum of 3 trials for each combination of variables was required for a neuron to be included in the analysis. A total of 1939 neurons were included. For dPCA analysis across performance levels, for each data point at a performance level *perf*, 200 neurons were bootstrapped from a pool of neurons recorded in sessions with performance ∈ [*perf* -5%, *perf* +5%], and repeated 100 times for the boxplot.

### Metacognitive behavioral model

To describe the mouse behavior, we employed a dual-agent approach, combining a rarity-based win-stay-lose-switch (r-WSLS) agent with a stimulus-guided reinforcement learning (RL) agent. Based on the observation that the choice updating WSLS strategy is stimulus-specific, stimulus A trials and stimulus B trials are separately processed in the model[7].

**Rarity-based WSLS agent.** This agent combines a WSLS-learning model[8] with a rarity state component. The WSLS-learning model allows update in the probabilities of staying and switching. Briefly, the probabilities of repeating a rewarded choice (win-stay) or to switch from an unrewarded choice (lose-switch) change over trials (t), according to:

$$p_{WS}(t+1) = p_{WS}(t) + \theta_{WS}\big(F_{WS} - p_{WS}(t)\big)$$
$$p_{LS}(t+1) = p_{LS}(t) + \theta_{LS}\big(F_{LS} - p_{LS}(t)\big)$$

In the formula above, $p_{WS}$ and $p_{LS}$ represent the probabilities to stay or switch, respectively; $\theta_{WS}$ and $\theta_{LS}$ represent the update rate of the probabilities; $F_{WS}$ and $F_{LS}$ represent the final, long-term values of the probabilities.

To computationally represent the internal rarity state, we used an unsigned prediction error as the basis for detecting a rare event[9]. We employed a simple value-based approach that is updated using the basic Rescorla-Wagner rule[10],

$$v_{s,a}(t+1) = v_{s,a}(t) + \gamma \cdot \varsigma_s(t) \cdot \Big(R(t) - v_{s,a}(t) \cdot \varsigma_s(t)\Big)$$

in which $s, a, R$ represents stimulus, action and outcome, $v_{s,a}(t)$ represents the recency-weighted value of a stimulus-action pair at given trial $t$, $\gamma$ represents the update rate, and $\varsigma_s(t)$ represents the saliency of stimulus $s$ at trial $t$. The unsigned discrepancy between actual and predicted value, *i.e.*, the absolute prediction error, is computed as

$$\Delta(t) = \big|R(t) - V_{s,a}(t) \cdot \varsigma_s(t)\big|$$

We incorporated an internal estimate of rarity, $\mathcal{R}$, that represents the agent's belief that the current trial is a common trial or a rare trial. For simplicity, we defined $\mathcal{R}_t$ to be a binary state value that is a Heaviside step function of the prediction error $\Delta(t)$ at trial $t$:

$$\mathcal{R}(t) = H(\Delta(t) - \mathcal{R}_{thresh})$$

in which $\mathcal{R}_{thresh}$ is the threshold to determine whether a trial is common or rare. We found this estimate sufficient to capture rare events, as it faithfully reflected most of the rare events in the trials.

Based on the behavioral data, we scaled the final probabilities $F_{WS}$ and $F_{LS}$ by the absolute prediction error $\Delta_t$ in each trial by the following empirical formula:

$$F_{WS}(t) = p_{perseverance} + \kappa_{WS} \cdot \Delta(\mathrm{t})$$
$$F_{LS}(t) = 1 - \big(p_{perseverance} + \kappa_{LS} \cdot \log(10^2 \cdot \Delta(t) + 1)\big)$$

To simulate the rarity-induced decision bias, we introduced an extra pair of parameters, representing the excessive probabilities of win-stay and lose-switch following rare+ and rare- trials. The probabilities are empirically defined as:

$$p_{\Delta WS} = \kappa_{\Delta \mathrm{WS}} \cdot \Delta(t) - \beta_{\Delta WS}$$
$$p_{\Delta LS} = \kappa_{\Delta LS} \cdot \Delta(t)^{\beta_{\Delta LS}}$$

The values of the coefficients $p_{perseverance}$, $\kappa_{WS}$, $\kappa_{LS}$, $\kappa_{\Delta \mathrm{WS}}$, $\beta_{\Delta WS}$, $\kappa_{\Delta LS}$, $\beta_{\Delta LS}$ are listed in Figure S9a, and are fixed for all simulations regardless of the inputs. To produce a choice in each trial, the r-WSLS agent tracks the last choice and outcome for each stimulus independently, and randomly choose to repeat the last choice (stay) or switch to the other choice (switch), depending on the last rarity state $\mathcal{R}(t-1)$: if the last trial is a common trial ($\mathcal{R}(t-1) = 0$), probabilities $p_{WS}$ and $p_{LS}$ are used; if the last trial is a rare trial ($\mathcal{R}(t-1) = 1$), probabilities $p_{WS} + p_{\Delta WS}$ and $p_{LS} + p_{\Delta LS}$ are used.

**Stimulus-guided reinforcement learning (RL) agent.** This agent employs a simple Q-Learning model with a multiplicative update rule[11] and distinct learning rates for positive and negative outcomes: if the choice corresponding to the stimulus is rewarded, the association between them ($s$ and $a$) is strengthened (with learning rate $\alpha_{R+}$), and the association between the stimulus and the opposite choice ($s$ and $a'$) is weakened (with learning rate $\alpha_{R-}$); conversely, if the trial was not rewarded, both associations are weakened (with learning rate $\alpha_{UR-}$)[7]. These rules are formulated as:

for rewarded trials,

$$\begin{cases} \delta_{s,a}(\mathrm{t}) = \alpha_{R+} \cdot Q_{s,a}(t) \cdot \left(R(t) - Q_{s,a}(t)\right) \\ \delta_{s,a'}(\mathrm{t}) = \alpha_{R-} \cdot Q_{s,a'}(t) \cdot \left(\epsilon - Q_{s,a'}(t)\right) \end{cases}$$

for unrewarded trials,

$$\delta_{s,*}(\mathrm{t}) = \alpha_{UR-} \cdot Q_{s,*}(t) \cdot \left(\epsilon - Q_{s,*}(t)\right)$$

In these formulae, $\delta_{s,a}(\mathrm{t})$ represents the update for stimulus-action pair $(s, a)$, $Q_{s,a}(t)$ represents the Q-value of the $(s, a)$ pair, and $\epsilon$ represents a relatively small noise value ($< 10^{-4}$). Choice from this agent is generated following a noisy epsilon-greedy approach:

$$p(\leftarrow) = N(Q_{s,\leftarrow} - Q_{s,\rightarrow}, \sigma^2)$$

**Entropy-based mixture-of-agents model.** To integrate the two agents to produce a unified metacognitive model, we defined normalized decision weights as the probabilities of following each agent's choices. Based on the recording data that: 1) rarity-modulated neurons were constantly present in the PPC, paralleling the persistence of the suboptimal rarity impact throughout learning; while 2) the stimulus-encoding capacity of PPC neurons increases, paralleling the performance improvement as learning progresses, we reasoned that the learning process is mainly driven by the RL agent. Thus, we employed an entropy-based metric[12] by transforming the Q-values into probability distributions using SoftMax function, computed the entropy over the distributions, and then used the entropy as the decision weight of the RL agent. Fixed decision weights were used for the r-WSLS agent before normalization. The final choice is randomly sampled from these choices with the decision weights as the probability distributions.

### Statistics

The normality of data was first examined using Lilliefors test. For normally distributed data, paired t-test or t-test were used for paired and unpaired data, respectively. For non-normal data, Wilcoxon signed-rank test or Wilcoxon ranksum test were used for paired and unpaired data, respectively. Statistical tests used in electrophysiological analyzes were described in detail in the "**Electrophysiological data analysis**" section.
